# Environment-dependent landscapes of coding variant impacts on coproporphyrinogen oxidase

**DOI:** 10.64898/2026.07.02.735896

**Authors:** Warren van Loggerenberg, Haotian Zhang, Vignesh Senguttuvan, Michael J. Chambers, Mailoan Panchalingam, Alireza Rasoulzadeh, Anna Axakova, Marinella Gebbia, Robert J. Desnick, Bruce Wang, Caroline Schmitt, Laurent Gouya, Jordi To-Figueras, Alexander Wahl, Ivet Bahar, Pemra Doruker, Frederick P. Roth

**Affiliations:** Donnelly Centre, University of Toronto, Toronto, Ontario, Canada; Department of Molecular Genetics, University of Toronto, Toronto, Ontario, Canada; Lunenfeld-Tanenbaum Research Institute, Sinai Health, Toronto, Ontario, Canada; Department of Computer Science, University of Toronto, Toronto, Ontario, Canada; Department of Computational and Systems Biology, School of Medicine, University of Pittsburgh, Pittsburgh, PA, USA; Department of Genetics and Genomic Sciences, Icahn School of Medicine at Mount Sinai, New York, NY, USA; Department of Medicine and Division of Gastroenterology, University of California San Francisco, San Francisco, CA, USA; French Centre of Porphyrias, AP-HP, Hôpital Louis Mourier, F-92700 Colombes, France; Université Paris Cité, INSERM, EFS, BIGR, F-75015 Paris, France; Biochemistry and Molecular Genetics Department, Hospital Clinic, IDIBAPS, University of Barcelona, Barcelona, Spain; Labcorp Genetics (Formerly Invitae Corp.), San Francisco, CA, USA; Laufer Center for Physical and Quantitative Biology and Department of Biochemistry and Cell Biology, School of Medicine, Stony Brook University, New York, NY, USA; Allotar Therapeutics, Pittsburgh, PA, USA

## Abstract

Hereditary coproporphyria (HCP) — caused by variants in coproporphyrinogen oxidase (*CPOX*) — can be diagnosed via genome sequencing. However, 74% of clinically-reported *CPOX* missense variants are classified as variants of uncertain significance (VUS) due to lack of evidence. *CPOX* variant classification is further complicated by environment-dependence: For example, the CPOX variant p.Asn272His (c.814A>C) is classified as benign yet has been associated with HCP-like symptoms in the context of mercury exposure. Here we measured the functional impact of nearly all possible CPOX amino acid substitutions in both the presence and absence of mercury. The resulting CPOX variant effect maps reflect known protein structure and mutational tolerance patterns while also offering new sequence-structure-function insights. Scores from this atlas not only distinguish pathogenic from benign variants but also identify mercury-dependent variant impacts, thus informing our clinical, structural, and functional understanding of CPOX deficiency and illustrating the value of systematic context-dependent multiplexed assays of genetic variant effects.

## 1. INTRODUCTION

Across genetic diseases, most clinically-encountered missense variants are classified as ‘Variants of Uncertain Significance’ (VUS) due to lack of evidence^1,2^. While results from (typically resource-intensive) validated functional assays can offer strong evidence^3^, they are unavailable for the vast majority of clinical variants. Recently, however, multiplexed assays of variant effect (MAVEs) have enabled measurement of functional impacts for nearly all substitutions in a target protein^4^, including those not yet observed clinically. The resulting variant effect maps can greatly reduce the ‘VUS problem’^4–6^. MAVEs are also beginning to systematically capture variant impacts that depend on genetic or environmental context ^7–10^.

The human coproporphyrinogen oxidase (*CPOX*) gene encodes CPOX (MIM: 612732, EC 1.3.3.3), the sixth enzyme in heme biosynthesis. CPOX deficiency causes hereditary coproporphyria (HCP; MIM: 121300), a rare autosomal dominant acute hepatic porphyria (AHP)^11^. Biochemical testing is often but not always definitive, whereas genetic testing can, in addition to confirming AHP, distinguish subtypes such as HCP, identify causal variants, and enable cascade screening of at-risk family members^12^.

Of the 244 clinical CPOX variants reported in ClinVar, 181 (74%) are annotated as VUS, with most (75%) of these being missense variants^13^. Further complicating interpretation, CPOX variant impacts can depend on the environment. For example, the common CPOX variant p.Asn272His (c.814A>C; global minor allele frequency 15%^14^) is classified as “benign” but nevertheless associated with an atypical porphyrinogenic response (APR) in people exposed to even low levels of Hg²⁺ (‘mercury’)^15,16^. APR is defined by the cascading upstream accumulation of porphyrin, including uroporphyrinogen III (URO; UROD’s substrate) and COPRO (UROD’s product and CPOX’s normal substrate) that results when mercury inhibits both CPOX and the immediately-upstream enzyme uroporphyrinogen decarboxylase (UROD; MIM: 613521, EC 4.1.1.37). Mercury sensitivity of the CPOX p.Asn272His variant arises from two factors: (1) mercury-induced accumulation of URO competes with the normal COPRO substrate; and (2) the CPOX p.Asn272His variant is catalytically promiscuous, more readily accepting URO as substrate and producing dehydroisocoproporphyrin, an atypical porphyrin that is subsequently converted to a ‘dead end’ metabolite (ketoisocoproporphyrin)^17^. Although p.Asn272His is the only currently-known CPOX variant subject to mercury-induced APR, there may be others.

Here, we measured the functional impacts of over 6,500 CPOX amino acid substitutions, both in the presence and absence of mercury. The resulting atlas of CPOX variant impact concords well with prior knowledge, while also offering sequence-structure-function insights. Moreover, this atlas represents a valuable resource for clinical understanding of *CPOX* variants, including those with mercury-dependence.

## 2. RESULTS

### 2.1 Scalable functional assays for CPOX missense variants

Using a humanized yeast model^18^ we confirmed that growth of yeast strain TSA1076 carrying a temperature-sensitive (ts) mutation in the essential *CPOX* ortholog *HEM13* could be rescued (at the non-permissive temperature) via expression of human *CPOX* cDNA (Figure S1A). In an initial evaluation using eight pathogenic and three benign missense variants, the assay showed the expected failure to rescue for seven (87.5%) of eight pathogenic variants and showed rescue for all three benign variants; Figure S1A). Among the variants confirmed to rescue (in the absence of mercury) was the ‘benign’ but mercury-dependent Asn272His variant noted above. Using the same assay, increasing mercury concentrations yielded reduced growth for both WT CPOX and the p.Asn272His variant, with the variant showing greater-than-WT sensitivity at intermediate mercury levels, consistent with its association with mercury-dependent APR (Figure S1B; see Methods).

### 2.2 Multiplexed assessment of *CPOX* variant effects

We next scaled this growth rescue assay using a competitively-grown pool of cells, each expressing a single *CPOX* variant clone, to obtain multiplexed measurements for all strains in parallel (Figure 1A; see Methods). Briefly, three mutagenic libraries were generated, with mutagenesis targeting a different third of the (full-length) coding sequence in each. Collectively encompassing nearly all amino acid substitutions (Figure S2D), each library was cloned into yeast expression vectors and transformed *en masse* into the *hem13*-ts strain. Each strain pool was grown competitively at the non-permissive temperature under both “baseline” and 96 µM mercury conditions. Functional impact scores were calculated from the log-ratio of post- to pre-selection variant frequencies^19^, and rescaled so that 0 and 1 corresponded to the median of nonsense and synonymous variants, respectively. The uncertainty (standard error) of each score was also estimated^7,20^. We thereby generated high-quality variant maps with functional impact scores in both the baseline and mercury context for >75% of all possible missense variants (88% of amino acid substitutions achievable via a single-nucleotide variant or SNV; Figure 1C; S2D).

**Figure 1.**
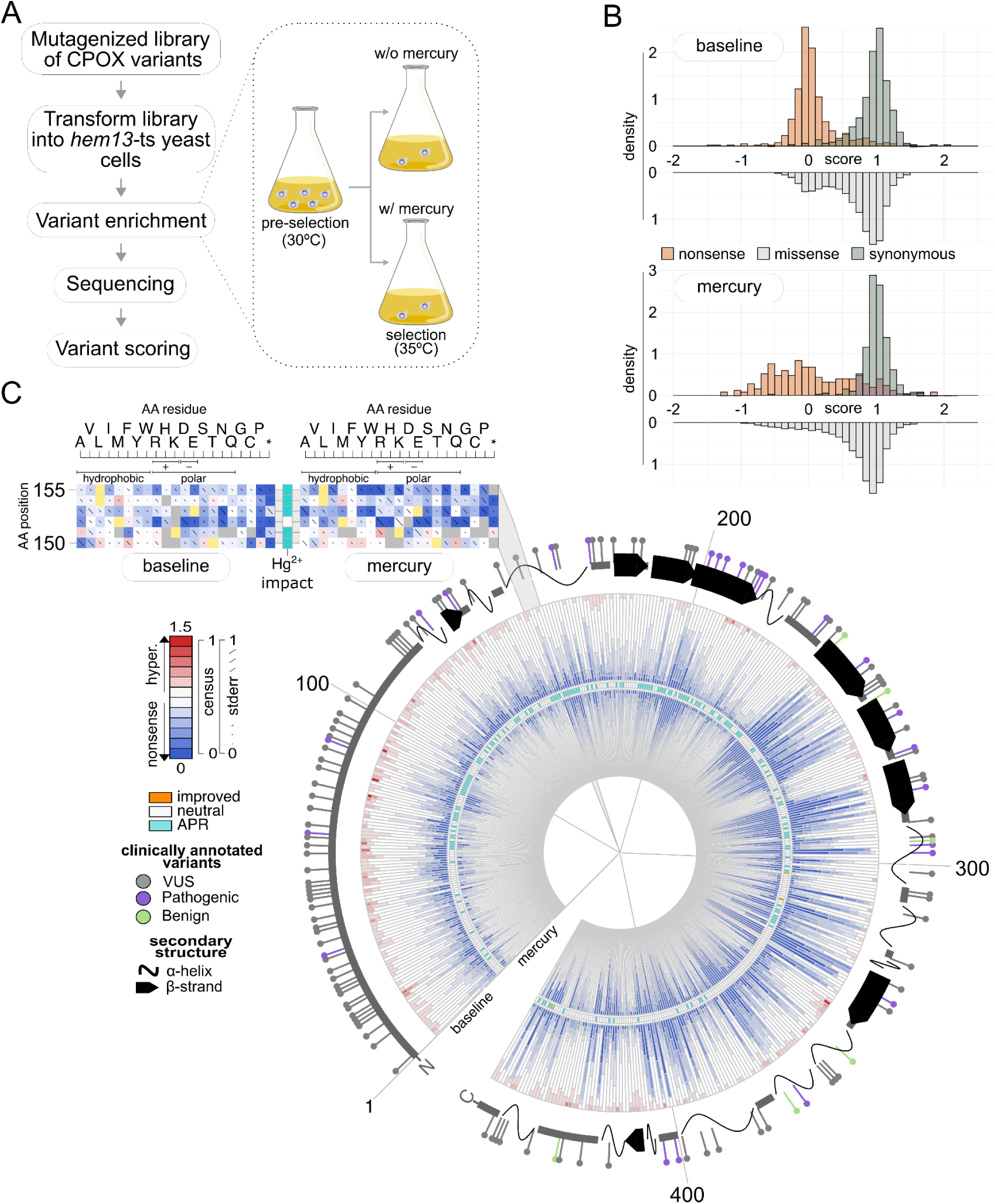
Mapping variant effects across CPOX with and without mercury. A. Overview of process to generate variant effect maps. B. Distributions of functional impact scores of nonsense (orange), synonymous (green), and missense variants (gray) under “baseline” conditions and in the presence of 96 µM mercury. C. Consensus tracks summarizing position-specific variant effects under baseline (outer ring) and mercury-treated (inner ring) conditions. Colors indicate damaging (blue), tolerated (white), or above-WT (“hyper”; red) substitutions. The central band highlights residues at which the majority of substitutions exhibited scores in baseline and mercury maps that together were consistent with an atypical porphyrinogenic response (APR; teal; see Methods). Predicted secondary structure is shown in the outermost ring (arrows, β-strands; loops, α-helices), with lollipops indicating clinically annotated benign (lime) and pathogenic (purple) variants. The expanded view of residues 150–155 exemplifies how a ‘wedge’ in circular representation corresponds to a segments from more traditional representations of the baseline and mercury variant effect maps (see Figure S16 for the full length versions). Here, each x-axis indicates amino acid substitutions, and each y-axis indicates CPOX residue position. Diagonal bars represent the estimated error for each variant effect score (0–1.5 scale).

To initially assess map quality, we examined the distributions of synonymous and nonsense variant scores and found them well separated for both maps (Figure 1B). Missense variants from each map showed a bimodal distribution (with modes that aligned well with nonsense and synonymous distributions), indicating that variants tended to either have a strong or neutral functional impact with relatively few variants having intermediate impacts. A small fraction (12%) of missense variants in both baseline and mercury maps appeared to “hyper-complement,” exhibiting growth beyond that of WT human CPOX in yeast. Although many of these fell within experimental error of the synonymous distribution, we investigated whether variants with apparently-increased fitness in our yeast assay tended to be deleterious in humans, as estimated based on depletion of such variants in non-human species as has been reported for other genes^20^. Indeed, this analysis applied to our baseline CPOX baseline map (see Methods; Supplementary Note 1) showed that hyper-complementing variants tend to be phylogenetically depleted and thus more likely to be deleterious.

As an additional quality control check, we sought to determine whether variants that appeared damaging in our baseline map tended to be counterselected in humans. Consistent with the observation that predicted loss-of-function stop and frameshift variants in CPOX are depleted (observed/expected ratio = 0.56)^14^, we found that CPOX missense variants with damaging baseline impact scores (see Material and Methods) tended to have lower allele frequencies (Figure S3A).

Impact scores were well correlated, both between biological replicates in each environment (Pearson’s *R* baseline = 0.86, *R* mercury = 0.85; Figure S2C) and between baseline and mercury maps (Pearson’s *R* = 0.73, Figure S3B). To specifically examine the impact of mercury on CPOX variant effects, we calculated a ‘delta’ score (mercury functional score − baseline functional score) for each variant (p<2e-16 by Wilcoxon test; see Methods). The distribution of delta scores was found to have a slight leftward skew (Figure S3C), suggesting that mercury reduces the enzymatic function of many variants.

We sought to compare our maps with previous experimental measurements of CPOX missense variant function, but the largest available set of consistently-collected CPOX missense variant function assay results was limited in two ways: 1) it encompassed only seven variants measured in both of our maps (Supplementary Dataset S11); and 2) these *in vitro* assays measured only specific (not total) activity. Given limited data, and the fact that our functional assay measured impact on total enzymatic activity (thus capturing not only impacts on specific activity but also those on protein abundance due to misfolding or degradation), it is unsurprising that correlation with both maps was not significant (P>0.1; Spearman Correlation Test; Figure S3E); However, scores from both maps did show modest but statistically-significant correlation with predicted stability impacts (baseline map Pearson’s *R =* 0.31, p <2e16; mercury map *R =* 0.32, p <2e16; Figure S3D).

### 2.3 Exploring the biochemical basis of CPOX variant effects

We next investigated trends related to physicochemical properties of amino acids. As expected, substitutions at hydrophobic residues were more damaging than those at polar (adjusted *p* [p_adj_] = 2e-15 and p_adj_ = 2e-8 for baseline and mercury maps, respectively; Wilcoxon test) or charged positions (mean score 0.87; p_adj_ = 6e-15 and 3e-6 for baseline and mercury maps, respectively; Figure S5A). Substitutions from hydrophobic to hydrophobic residues were generally tolerated (mean score 0.85), whereas substitutions to proline—with a known tendency to disrupt secondary structure—were generally less tolerated (mean score 0.59; p_adj_ <8e-22 and p_adj_ = 4e-6 for baseline and mercury maps respectively; Wilcoxon test; Figure S5B, S5C).

We further probed variant impacts in the context of CPOX’s monomeric and homodimeric structures. As expected, we found buried residues (here defined by solvent-accessible surface area [ASA] <25%) to have a lower median score than surface residues not at the dimer interface (>50% ASA; Δmedian = 0.2; p = 5e-16, Δmedian = 0.25; p = 1e-14, for baseline and mercury map, respectively; Figure S6B). Interfacial residues similarly exhibited significantly lower scores than non-homodimer surface residues (Δmedian = 0.16; p = 1e-3, Δmedian = 0.16; p = 3e-4, for baseline and mercury map; Figure S6B), consistent with the known stabilizing role of CPOX dimerization^21^.

Examining our maps in the context of the mitochondrial targeting sequence (MTS; residues 1–110)^11^, we found substitutions generally tolerated across characteristic MTS features^22^. However, substitutions in the hydrophobic stretch of the intermembrane space targeting (IST) domain (residues 75–93) were poorly tolerated in the baseline (Δmedian = 0.1 and p = 1e−2; Wilcoxon test; Figure S6C) but not the mercury map (Δmedian = 0.01, p = 0.3; Wilcoxon test). Human CPOX (unlike the cytosolic yeast Hem13 ortholog) is targeted to the intermembrane space, where it must be ‘peripherally attached’ to the inner mitochondrial membrane^23,24^. While our baseline map is consistent with failure of IST domain variants to peripherally attach, it is unclear how mercury binding would diminish these effects (discussed further below).

To begin evaluating our maps in the context of CPOX’s ability to bind substrate (Figure 2A), we examined Arg262, Arg328, Arg332, and Arg389. The Arg262 residue mediates recognition of COPRO, upon which CPOX carries out (metal- and cofactor-independent) oxidative decarboxylation of COPRO’s propionic acid side chains to form protoporphyrinogen IX^21^, Arg332 displayed intolerance while Arg328 and Arg389 each exhibited tolerance to substitution (with maps showing concordance for all three), suggesting lower importance for Arg328 and Arg389.

**Figure 2.**
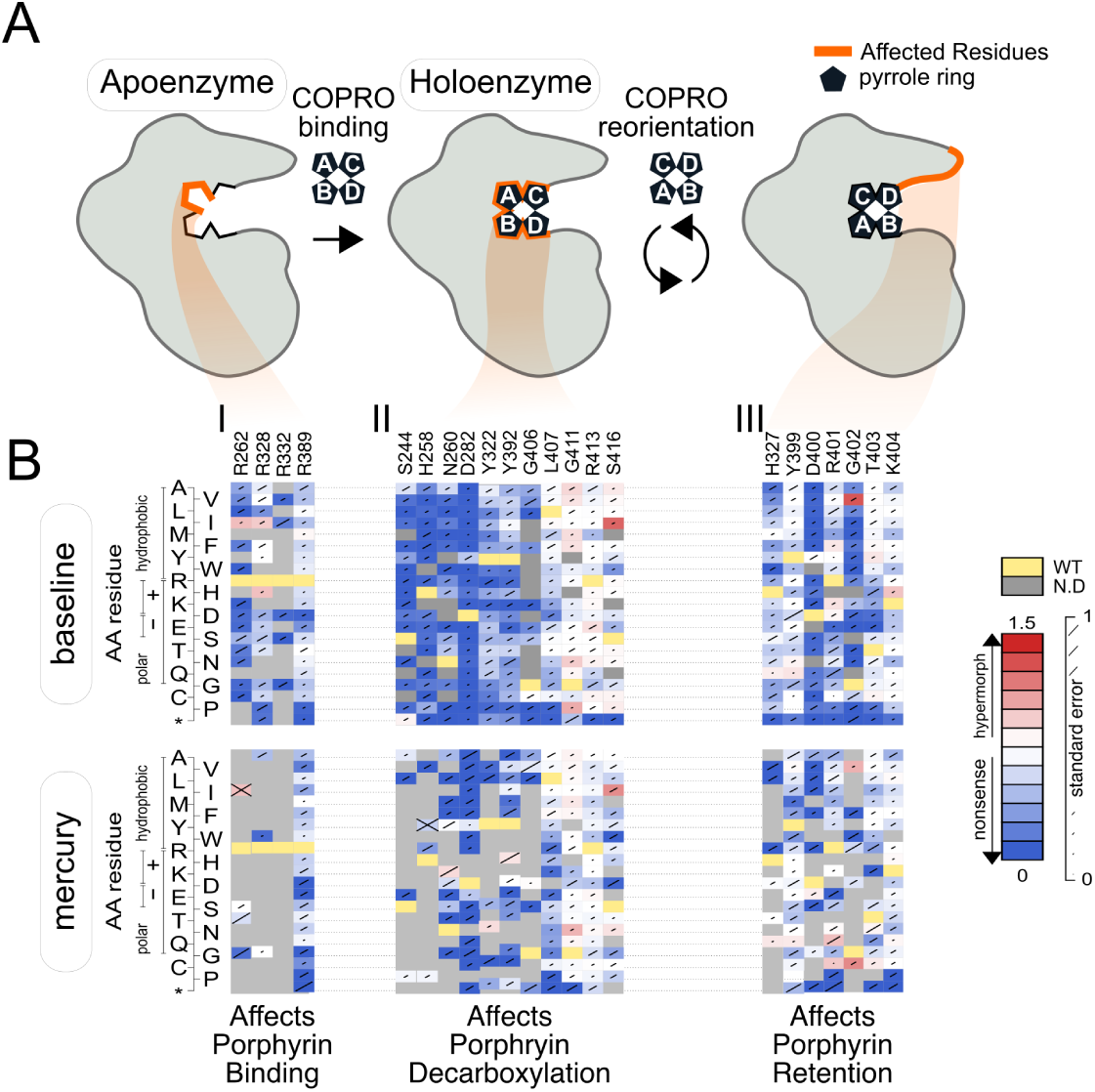
Map scores reveal patterns of mutational tolerance in the active site. A. A schematic of CPOX’s two-step oxidative decarboxylation of coproporphyrinogen III (COPRO), involving constrained orientation of its intermediate substrate. B. Baseline and mercury map scores for each possible substituted amino acid (y axis) at each active site residue position (x axis) involved in: (I) mediating substrate recognition, (II) catalysis, and (III) deterring premature release of its intermediate substrate. For each substitution, diagonal bar sizes convey estimated measurement error in the corresponding score. Box color either indicates the WT residue (yellow); a substitution with damaging (blue), tolerated (white), or above-WT (“hyper”, red) functional score; or not measured (gray).

Next, we examined residues related to enzyme kinetics. Both His258 and Ser244, each important for enzyme kinetics, were intolerant to substitution in our maps as expected^21^ (Asp282, Asn260, Ser416, Arg413, Gly411, Leu407, Gly406), our maps found that only Asp282, Asn260, and Gly406 were generally intolerant to variation (Figure 2B, II), suggesting that Ser416, Arg413, Gly411, and Leu407 are dispensable for this function.

We next examined the active site loop (residues 400-413; Figure 2B, III) thought to be important for retention of harderoporphyrinogen (formed via decarboxylation of COPRO) and its distortion to favor decarboxylation^21^. More specifically, we examined residues Asp400-Lys404 for which some variants are known to prematurely release the harderoporphyrinogen intermediate instead of converting it to PPIX^25^. Such variants, when homozygous, are associated with Harderoporphyria (HP; also known as HARPO), a clinically distinct variant of AHP^26^. Among this Asp400-Lys404 subset of active site loop residues, our data only implicated Asp400 and Gly402 as being critical for CPOX function. We were initially surprised to find Arg401 and Lys404 generally tolerant to mutations, as Arg401 is highly conserved and p.Lys404Glu is a known pathogenic variant. However, this *in vitro* tolerance is consistent with the observation that individuals homozygous for p.Lys404Glu, despite reduced ability to convert harderoporphyrinogen to PPIX, do not experience acute HCP attacks^27^.

Although substitutions that newly introduce a negatively charged amino acid were generally less well tolerated than other substitutions (Δmedian = 0.08; p_adj_ = 4e-4; Wilcoxon test; Figure S5B, S5C), introducing negatively charged amino acids at residues Asp400-Lys404 tended to be even less well tolerated (Δmedian = 0.76 and p = 7e-3 for the baseline map; Wilcoxon test; Figure S7A). A role for electropositivity in Asp400-Lys404 fits the previously reported role of solvent-exposed electropositive residues in the active site (Arg413, Arg262, and His258), which engage with propionyl groups of the COPRO substrate^21^), rather than in substrate recognition or in catalyzing decarboxylation directly.

The strong correlation between our maps noted above includes correlation at residues near the substrate. For example, in the context of decarboxylation corridor residues facilitating substrate passage to the active site (Ser416, Arg413, Gly411, and Leu407), introducing negatively charged residues did not show a striking impact (Δmedian = 0.03 and p = 5e-1 for the baseline map; Δmedian = 0.25 and p = 2e-1 for the mercury map; Wilcoxon test; Figure S7B). An interaction between the residue p.His327 (helix α7), for which the p.His327Arg variant is associated with the Harderoporphyria phenotype, and Tyr399 (helix α9) was also predicted to play a key role in positioning the active site loop^27^; however, neither map implicated these residues as essential for function (Figure 2). For other substrate-proximal residues, we observed striking differences between the maps: The damaging effect of introducing negatively charged amino acids to active site loop residues Asp400-Lys404, which had been observed in the baseline map, was not observed in the mercury map (Δmedian = 0.07 and p = 0.4; Wilcoxon test; Figure S7A). One potential explanation may arise from the fact that mercury broadly suppresses CPOX’s second decarboxylation of harderoporphyrinogen^28^, so that substitutions that inhibit the second decarboxylation step yield ‘diminishing returns’ in terms of causing further harm, thus appearing less damaging under mercury conditions than do substitutions that reduce activity of the first decarboxylation step.

### 2.4 Prioritizing variants and structures to investigate mercury-sensitive CPOX missense variants

Mercury binding to CPOX (via the thiol groups of Cys319) has been suggested to allosterically alter substrate affinity, impairing the use of harderoporphyrinogen^28^. However, cysteine residues (including Cys319, Cys281 and Cys357) showed concordant effects between maps, suggesting that mercury-sensitive variant effects are not solely driven by altered cysteine accessibility (see Supplementary Note 2E). Indeed, analysis of ‘delta’ (mercury − baseline) scores showed that mercury reduced function for the majority (71%) of variants, but for only 41% of variants at these cysteine positions.

We sought to identify candidate APR-causing CPOX variants more broadly by identifying (amongst variants tolerated in the baseline map) missense variants with a significantly lower delta score than control variants (n = 2637, 43%; see Methods). Importantly, these candidates included the only well-established mercury-dependent APR variant p.Asn272His as well as p.Val135Ala, observed previously in an individual diagnosed with HCP^29^ after suspected heavy-metal exposure (observed clinically by author J. To-Figueras; Figure S16). To obtain a more confident subset of candidates, we next considered the 151 positions where the majority of substitutions were candidates in the initial set (shown in teal in the band between baseline and mercury maps in Figure 1C), and this refined set included 1403 variants (with p.Asn272His and p.Val135Ala still among them).

How CPOX p.Asn272His confers mercury-dependent preference for the URO substrate remains unclear. Hypothesizing that this might arise via allosteric communication to the active site, we sought sequence-structure-function trends amongst our candidate mercury-sensitive substitutions. Interestingly, the refined candidate set was enriched for variants in α-helix 3 (α3; residues 147-160) and in β strands 1-3 (residues 183-213; Figure S8B). As neither of the presumed mercury-sensitive variants p.Asn272His and p.Val135Ala fall within these regions, the putative association of these secondary structural features with APR could not have been made purely based on the genotypes of patients with APR. To identify residues within APR-implicated secondary structures that are critical for functional dynamics and allosteric communication, we analyzed global modes of motion and found that Leu155 (α3) and Gly188 (β2) act as key hinge residues (see Figure S11F,G)^30^. Below we describe results from MD simulations on a set of representative mercury-sensitive substitutions (the ‘MSS-MD set’) (one at each of the four positions): p.Leu155Trp (α3; Figure 1D), p.Gly188Gln (β2), and p.Val135Ala (loop between α1–α2). p.Asn272His and WT were included as positive and negative controls, respectively (Table S2), as well as results from separate docking simulations of the COPRO and URO substrates.

First, however, we needed to select the appropriate structure(s) for which to model dynamics. The AlphaFold3 (AF) model and the crystal structure^21^ of human CPOX differ primarily in the conformation of the active-site loop (α10): it forms a helix in the AF model but appears unstructured in the human crystal structure (Figure S9). Notably, our baseline map shows proline substitutions within α10 to be deleterious, consistent with the structured helix of the AF model. Therefore, we present MD simulation results using the AF model, but reached similar conclusions using the crystal structure (see Supplementary Material).

#### 2.4.1 Impact of CPOX MSS-MD variants on protein secondary structure

Simulations showed that hinge position variants p.Leu155Trp and p.Gly188Gln allosterically affected the local conformational preferences of active site residues (i.e. Thr256, Arg262 and Tyr263) on β5 (Figure 3C). Thr256 fell within an extended (β-sheet) conformation in nearly all (98%) of WT snapshots, whereas within the p.Gly188Gln structure, Thr256 fell within a coil for 18% of snapshots, which suggests unfolding of β5. Similarly, in the p.Leu155Trp structure, Arg262 and Tyr263 showed reduced β-sheet conformation (58% down from 100% in the WT structure). These results point to allosteric effects affecting substrate preference at the active site as an explanation for the appearance of Thr256 and Tyr263 as APR candidates from our maps.

**Figure 3.**
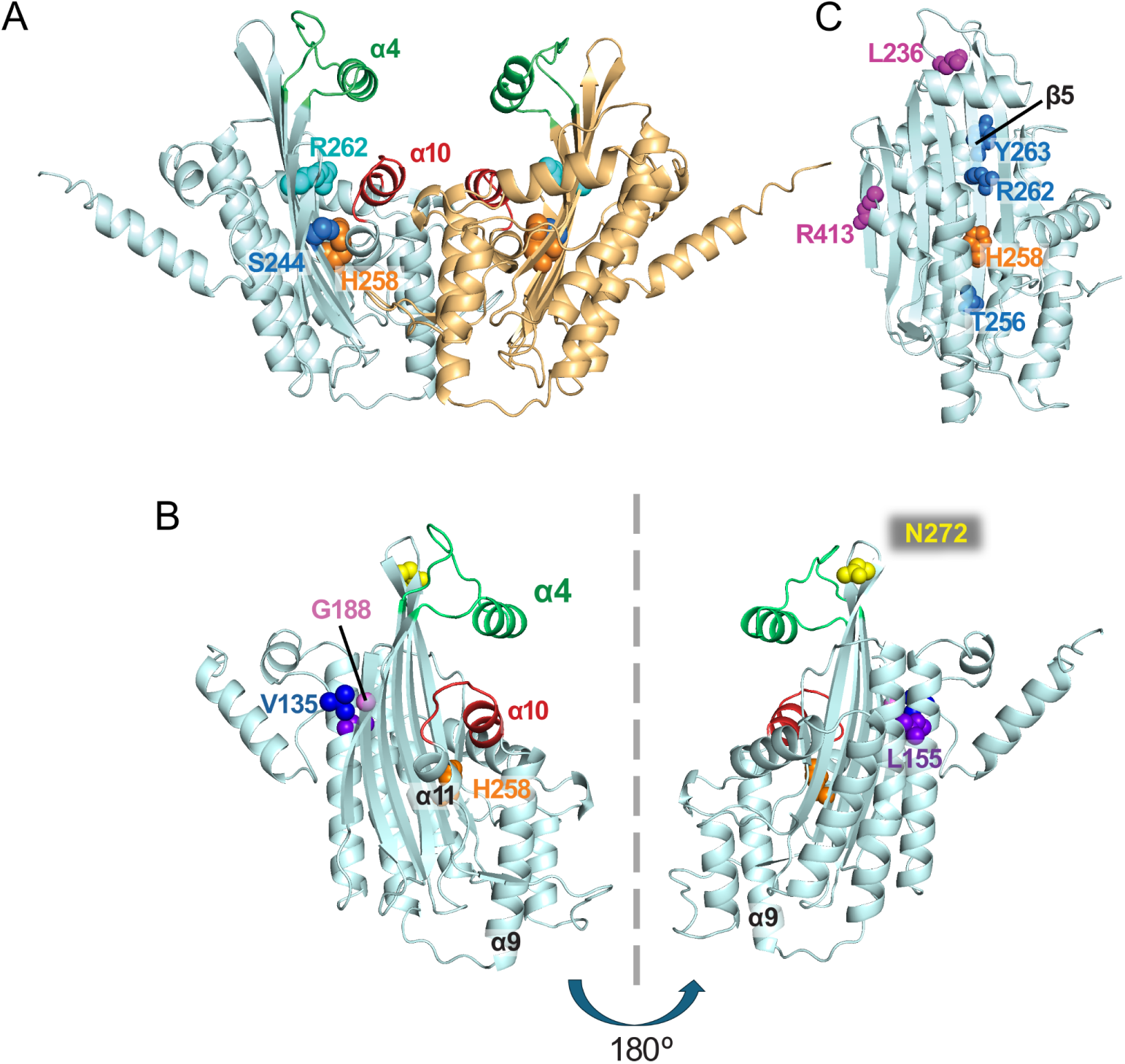
Structural details of human CPOX (AF model). **(A)** In each chain of the CPOX homodimer, the active site lies between the β-sheet and helices α9-α11. The main entrance to the active site (gate 1) is controlled by helices α10 on the active site loop (red) and α4 on the lid (green). The catalytic residue H258 (on β5) and key residues involved in ligand recognition (Arg262 on β5, Ser244 on β4) are represented with spheres. **(B)** Residues corresponding to the missense variants analyzed in our simulations (Asn272, Leu155, Val135 and Gly188) are shown from two views of a single chain. **(C)** The p.Gly188Gln and p.Leu155Trp variants allosterically induce local unfolding of specific residues near the active site (blue spheres on β5). In contrast, the p.Asn272His and p.Val135Ala variants alter local conformational preferences of Leu236 and Arg413 residues (magenta), respectively.

We also observed subtle conformational differences for variants, with potential to influence lid and active-site loop dynamics (Figure 3C). Specifically, Leu236 on the lid (adjacent to β4) fell within an extended (β-sheet) conformation in 31% of WT snapshots, decreasing to 19% in the p.Asn272His structure. Arg413 on the active-site loop (adjacent to α11) fell within an α-helical conformation in 11% of WT snapshots, but was never observed to be α-helical in the p.Val135Ala structure. These observations suggest that p.Val135Ala exerts a long-range allosteric effect on the active site, providing a potential explanation for the unusually severe clinical manifestations reported in two independent HCP cases for which p.Val135Ala was observed ^29,31^.

#### 2.4.2 Functional dynamics of WT CPOX vs. MSS-MD variants

##### Global motions

To capture the dominant modes of CPOX dynamics, we performed principal component analysis (PCA) of the trajectories generated by MD simulations. Examining the first two principal components (PC1 and PC2; Figure 4A,B), PC1 revealed a hinge motion between the two monomers that opened and closed via anti-correlated movements of the lid and active-site loop, while the motions along PC2 were concertedcounter-rotations of the two monomers. Conformational preferences of WT CPOX and its variants were projected onto the PC1-PC2 space as density maps (Figure 4C). While WT conformations were tightly clustered, variants exhibited broader conformational distributions.

**Figure 4.**
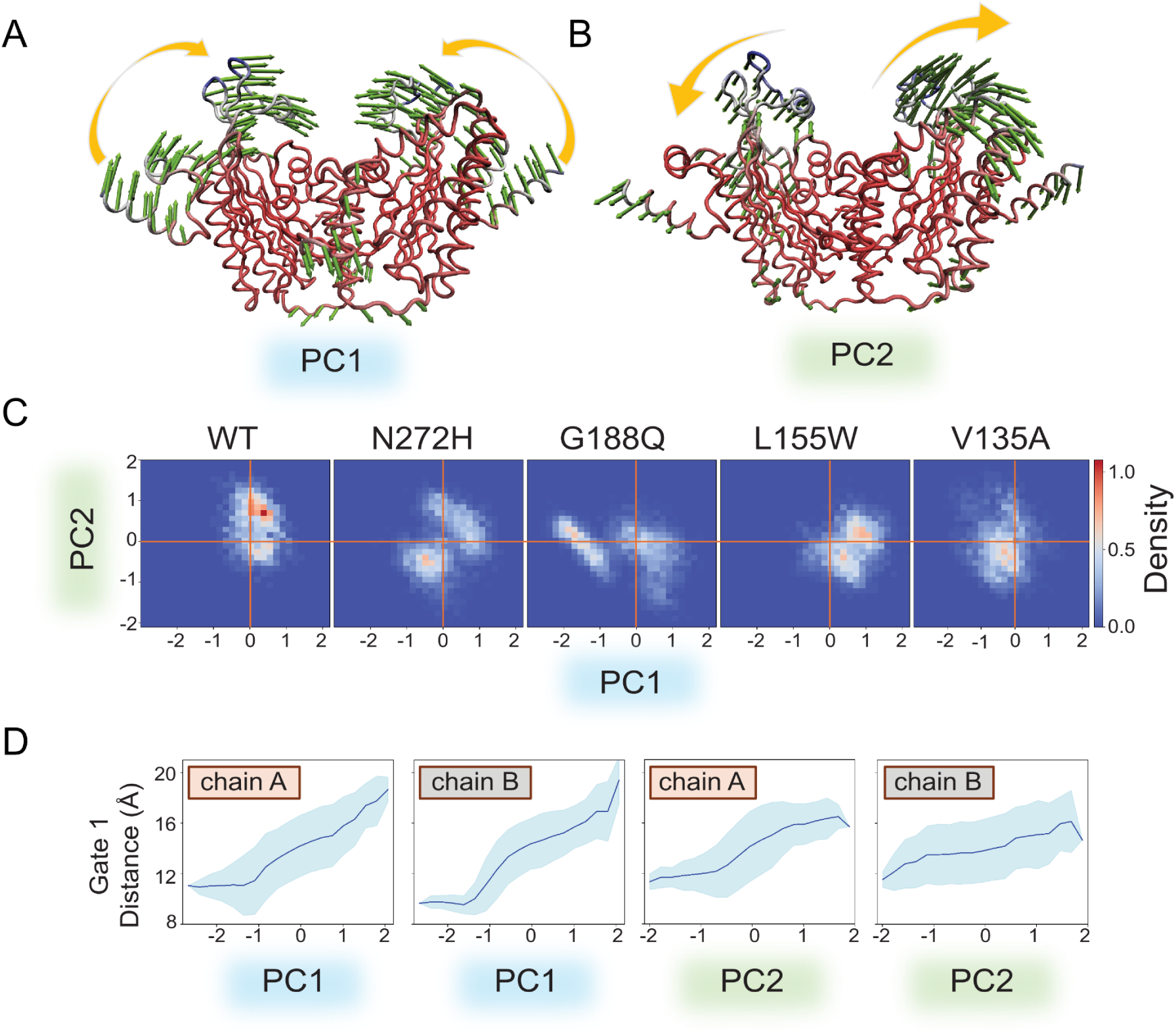
Global motions of CPOX (AF model), obtained by principal component (PC) analysis performed on all MD trajectories after structural alignment. **(A)** PC1 and **(B)** PC2 illustrate the deformation vectors (green) describing the global structural changes that are functionally relevant. **(C)** Population density maps plotted for WT and each variant, projected along the dominant directions PC1 and PC2. D) Gate 1 opening mediated by the first two principal components (PC1 and PC2).

##### Substrate entrance

We identified two primary entrances (gates 1 and 2) to the active site pocket (Figure 5A), and monitored their opening using the distance between key residue pairs (Table S3; Figure 5B). Gate 1 is regulated by the relative motions of helices α4 and α10, which has been observed in yeast structures (Figure S9E,F)^32^. Gate 1 remained open in about half of WT snapshots (distance >15 Å), and in a reduced but still substantial fraction of snapshots for all MSS-MD variants. This opening is largely driven by the bending motion captured by PC1 (Figure 4D). Conformational changes along PC2 also contributed to Gate 1 opening (Figure 4D). Gate 2, adjacent to Gate 1 and controlled by motions of α10 and β7, was less frequently open but did show some variation along both PC1 and PC2 (Figure S14). When Gate 2 opened, this was often in concert with Gate 1 as a composite opening, but was also the sole active entrance site in some snapshots.

**Figure 5.**
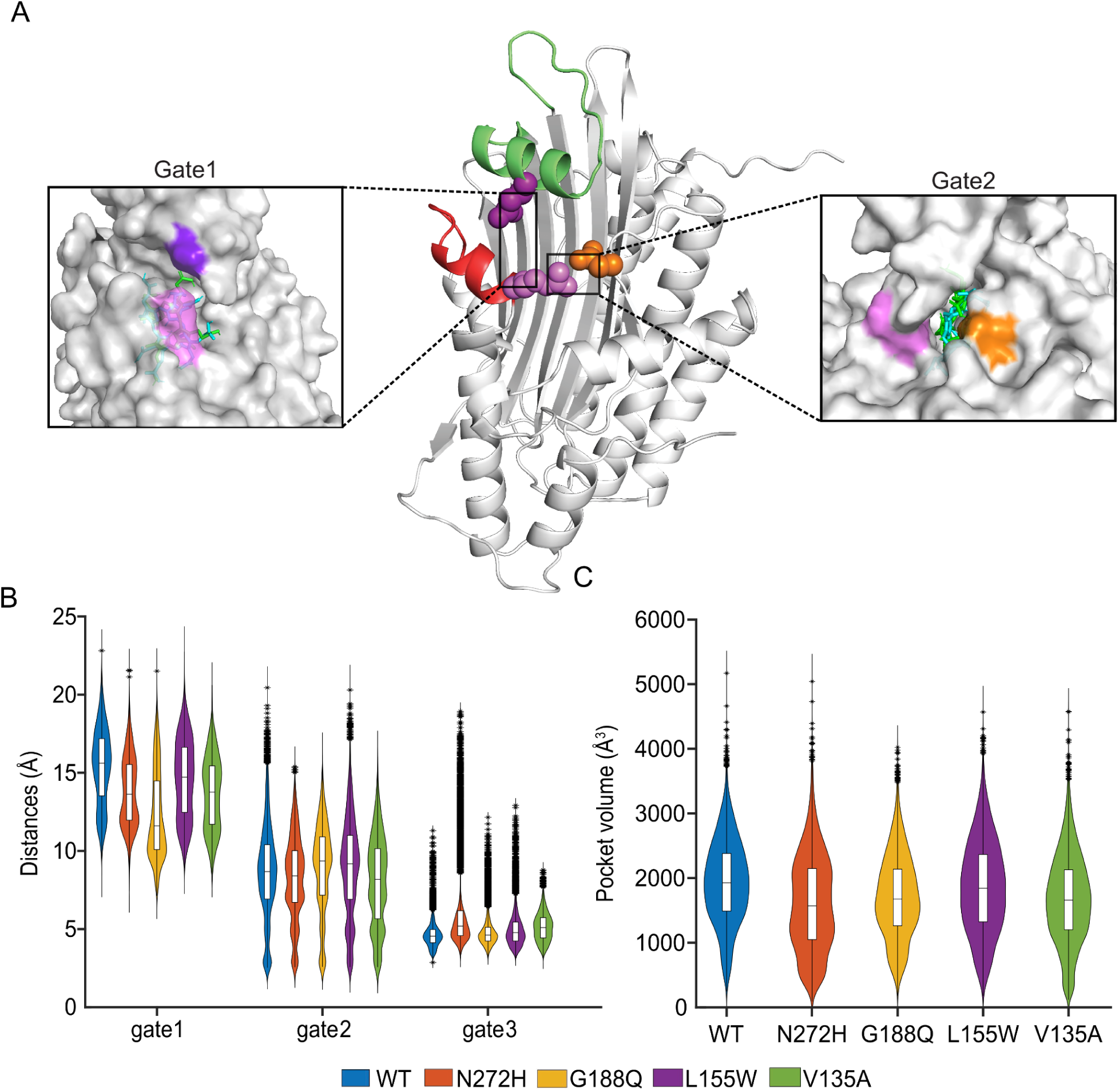
Characterizing active site entrances and pockets in molecular dynamics simulations (based on the AF structure model) of WT and variants. **(A)** Residues used to monitor the opening of primary gates 1 and 2 are shown as colored spheres (middle panel). The left box shows the surface representation of a snapshot with gate 1 open, highlighting the residues Gln221 (dark purple) and Arg401 (light purple). The distance between these residues is used for measuring gate 1 opening (see Table S3).Docked substrates COPRO (green) and URO (cyan) are shown as sticks within the active pocket. The right box shows another snapshot with gate 2 open and docked ligands; gate 2 distance is calculated between residues Asp340 (orange) and Arg401 (light purple). **(B)** Distance distributions for gates 1 and 2 are displayed as box plots for WT and variant runs. An additional gate 3 only opens in p.Asn272His. (**C**) Distributions of active site pocket volumes show that WT pockets are slightly larger on average than those of the variants.

##### Active site pockets

Pocket volume analysis for both monomers showed broad distributions of active site pockets’ sizes in WT and MSS-MD variants, with median values exceeding 1600 Å³ (Figure 5C). Pockets in the WT structure tended to be slightly larger than those in MSS-MD variant structures. Given the size of COPRO and URO, substrate entry requires not only a sufficiently wide opening but also adequate pocket depth to reach the catalytic site. While pocket volumes and gate distances can suggest open conformations, full substrate accessibility may also depend on side-chain rearrangements.

#### 2.4.3 Further interrogating impact on substrate preferences via docking

To explore binding preferences for COPRO and URO, we used snapshots judged to offer the best chance of productive substrate docking, as a starting point for simulated docking. Among snapshots with at least one open gate (distance > 15 Å or > 10 Å for Gate 1 and 2, respectively), we selected those with the largest pocket volumes. After five rounds of docking, we derived the active site docking success rate as the fraction of poses where catalytic residue His258 contacted the substrate.

Using the WT structure, the docking success rate was similar (∼60%) for both substrates. In contrast, across the four MSS-MD variant structures, success rates for COPRO ranged from 35-68% while for URO it was 61-90%, with URO’s increase in absolute success rate ranging from 21-52%. While docking does not explicitly account for entropy changes, these results suggest that MSS-MD variants have increased preference for URO.

Docking also enabled predictions of binding energy. The WT structure showed stronger binding to COPRO (binding energy −11.1 kcal/mol) than URO (−10.3 kcal/mol; ΔCOPRO-URO = -0.8 kcal/mol). This preference shifted slightly (towards stronger binding for URO) for the MSS-MD variants p.Asn272His (ΔCOPRO-URO = -0.3 kcal/mol) and p.Leu155Trp (ΔCOPRO-URO =-0.1kcal/mol). The preference shift towards URO binding was stronger for MSS-MD variants p.Gly188Gln (ΔCOPRO-URO=0 kcal/mol and p.Val135Ala (ΔCOPRO-URO=0.8 kcal/mol; see details in Table 1).

**Table 1.** Substrate docking using snapshots with helical active site loop (AF model)

| System | Docking results |  |  |  |  |  | MD simulations |  |
| --- | --- | --- | --- | --- | --- | --- | --- | --- |
|  | Binding affinity for best pose (kcal/mol) |  |  | Active site occupancy (%) |  | Total number of poses | Binding affinity from PRODIGY-LIG (kcal/mol) |  |
| | COPRO | URO | $\Delta$(COPRO-URO) | COPRO | URO | | COPRO | URO |
| WT | -11.1 | -10.3 | -0.8 | 59 | 60 | 300 | -12.5 ± 0.3 | -12.6 ± 0.4 |
| N272H | -10.8 | -10.5 | -0.3 | 68 | 89 | 200 | -11.6 ± 0.3 | -13.1 ± 0.3 |
| L155W | -10.8 | -10.7 | -0.1 | 35 | 61 | 200 | -11.7 ± 0.4 | -13.2 ± 0.4 |
| G188Q | -10.1 | -10.1 | 0.0 | 38 | 90 | 200 | -12.5 ± 0.4 | -13.2 ± 0.4 |
| V135A | -10.4 | -11.2 | 0.8 | 46 | 81 | 200 | -12.6 ± 0.2 | -13.5 ± 0.2 |

Subsequent MD simulations initiated from best docked poses showed that both substrates remained stably within the active site. PRODIGY-LIG affinity estimates^33^ also indicated stronger binding for URO than COPRO for the MSS-MD variants (Table 1, Supplementary Notes 2D). Taken together with the docking results, these results suggest that WT CPOX preferentially binds COPRO, whereas MSS-MD variants favor URO over COPRO.

### 2.5 CPOX variant effect maps distinguish pathogenic from benign variation

To assess the utility of our baseline map scores in distinguishing pathogenic from benign variants, we assembled two reference sets of missense variants: 1) a “positive” set of pathogenic or likely pathogenic (19 HCP- and 6 HP-associated) variants; and 2) a “negative” set with 7 missense variants classified as benign or likely benign (see Methods). Supporting our map’s quality, scores tended to be lower for positive than negative reference variants (Δmedian = 0.32; p = 1e-2; Mann-Whitney U test; Figure 6A). To further assess performance, we calculated both precision—the proportion of variants below a threshold score that are positive—and recall, the proportion of all positive variants that scored below the threshold. Because the fraction of all reference variants that fell in the positive set is unlikely to match the true prior probability of pathogenicity, we derived from each precision estimate the ‘balanced precision’ expected given a uniform (50%) prior probability of pathogenicity. Performance measures were: 1) area under the balanced precision recall curve (AUBPRC) and 2) recall at a stringent threshold of 90% balanced precision (R90BP). Both maps outperformed the average AUBPRC of 0.56 and R90BP of 0% observed from 1000 sets of randomly-permuted predictions: The mercury map yielded AUBPRC=0.85 and R90BP=52%, while the baseline map performed best with AUBPRC=0.89 and R90BP=62% (Figure 6B).

**Figure 6.**
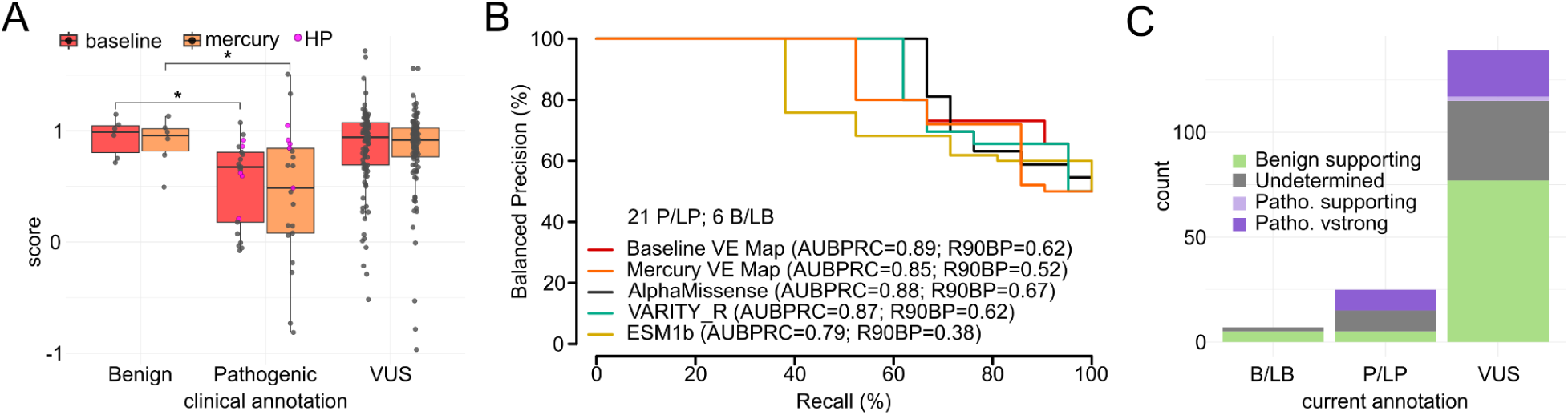
Variant effect maps accurately identify pathogenic variants and provide evidence for VUS reclassification. A. The distribution of functional impact scores from baseline (red) and mercury (orange) maps for reference “pathogenic,” “benign,” and “VUS” variant sets. Variants associated with harderoporphyria (lime) were also plotted. Boxes correspond to interquartile range, and bold bars indicate medians. Whiskers correspond to minima and maxima. Significance was evaluated with a Mann-Whitney U test. B. Evaluation of precision (fraction of variant scoring below each threshold functional score that are in the positive reference set containing pathogenic variants) vs. recall (fraction of positive reference variants with functional scores below threshold). Here, precision values have been “balanced” to reflect performance in a setting where positive and negative sets contain the same number of variants. Balanced precision-recall curves are shown for baseline (red) and mercury maps (orange), as well as the computational predictors AlphaMissense (black), VARITY (turquoise) and ESM1b (gold). Performance is also described in terms of area under the balanced precision vs. recall curve (AUBPRC) and recall at a balanced precision of 90% (R90BP). C. Currently clinically annotated variants and new evidence proposed based on calibrating our baseline map scores against the reference set (see Methods for details on the derivation of evidence strength).

Because computational variant effect predictors (VEPs) are an increasingly useful source of evidence about variant pathogenicity^34–37^, we compared map performance with VEPs. We first note that, according to American College of Medical Genetics and Genomics and Association for Molecular Pathology (ACMG/AMP) guidelines^3^, VEP and functional assay evidence are not in competition, as they are considered to be independent evidence sources. Nevertheless, we benchmarked the VEPs AlphaMissense^38^ (AUBPRC = 0.88; R90BP = 67%), VARITY^39^ (AUBPRC = 0.87; R90BP = 62%), and ESM-1b^40^ (AUBPRC = 0.79; R90BP = 38%, Figure 6B), finding the best of these to be roughly on par with the performance of our baseline map.

Given that human CPOX can rescue the (cytoplasmic) HEM13 ortholog in yeast cells, our assay was unlikely *a priori* to identify defects in the MTS. However, excluding the MTS positions had minimal effect on either map’s performance (R90BP = 61% and 56%, respectively), whereas excluding these positions modestly improved VEP performance (R90BP = 78%, 72%, and 44%, respectively; an increase of 5-6%; Figure S15B).

The baseline map (using a stringent threshold achieving 90% balanced-precision) identified three of five known HP-associated pathogenic variants (including p.Gln162Pro), while the mercury map with a similarly stringent threshold identified only p.Gln162Pro. When HP-related positions were excluded, recall improved for the mercury map (R90BP = 62% vs 52% previously), was unchanged for the baseline map, and was lower by 5-7% for VEPs (R90BP = 62%, 56%, and 31%, respectively; Figure S15D). While our maps (especially the mercury map) will not be highly useful as evidence towards benignity for the distinct HP subtype of AHP, they should prove useful in identifying additional pathogenic HP variants.

Finally, to calibrate the evidence strength that our measurements can provide towards variant pathogenicity classification, we applied an established kernel density estimation method to our reference sets and derived a likelihood ratio of pathogenicity (LLRp; Figure S15E; see Methods) for each variant score^41,42^. LLRp values can in turn be calibrated to evidence strength levels within the ACMG/AMP framework^20,42^. Thus, our maps offered new functional evidence for 101 (73%) of the 136 CPOX missense VUSs reported in ClinVar (Figure 6C; see Supplementary Dataset S4). Of these 101 variants, 77 (76%) received supporting evidence towards benignity, while 2 (2%) and 22 (22%) received either supporting or very strong evidence, respectively, towards pathogenicity.

## 3. DISCUSSION

Here we systematically assessed CPOX variant functions with and without mercury, which can increase the penetrance of *CPOX* variants. By combining comprehensive codon mutagenesis with large-scale multiplexed functional assays, we experimentally measured the impacts of nearly all possible *CPOX* missense variants, revealing environment-dependent impact changes for >40% of missense variants.

Both maps recapitulated many known sequence–structure–function relationships. Buried positions scored significantly lower in both maps, as expected. Missense variants at COPRO-binding positions and in the decarboxylation corridor were generally damaging. Residues closer to the active site tended to be less tolerant of substitution. The active-site loop was largely intolerant to introduction of negative charge, likely because this disrupts retention of negatively charged harderoporphyrinogen after substrate re-orientation ahead of the second decarboxylation.^21^.

Mercury inhibited both WT and variant CPOX proteins, consistent with a previous report^28^. Simulations showed that p.Asn272His and p.Val135Ala (known and suspected, respectively) residues causing mercury-induced APR and allosteric impacts on CPOX enzyme kinetics perturb active site loop and lid dynamics. Our maps identified additional such residues: For example, MD simulations showed p.Gly188Gln and p.Leu155Trp locally destabilized β5 residues forming the base of the active site, altering pocket volumes and gate distances critical for substrate entry, with global chain bending and anticorrelated lid movements that further modulated active site accessibility. Indeed, all four mercury-sensitive variants we simulated showed not-seen-in-WT conformational states favoring URO over COPRO substrates, consistent with mercury-induced APR caused by altered substrate handling.

Our simulations had limitations. While they clearly showed the helical conformation to be critical for substrate binding and catalysis, they may have missed rare or slow transitions such as those between unstructured and helical states of the active site loop. Also, simulations have limited efficiency to capture binding events for large substrates such as COPRO and URO. However, the latter was addressed, at least in part, by complementing simulations with docking analysis for a subset of ‘MD snapshots’.

Our experimental assays also had limitations. One caveat is that experimental measurements are subject to random error, estimated here for each variant using established methods^19^. Systematic errors are also possible: For example, we expressed variants only in the context of a mature cDNA and so inevitably missed not only non-coding variants but also the impact of coding region variants on splicing. Also, C-terminal nonsense variants which appeared tolerated in our assay may nevertheless trigger nonsense-mediated decay in the endogenous context. Also, our assay was yeast-based, measuring rescue of the cytosolic yeast ortholog Hem13^23^, so that changes in the human CPOX’s MTS, which were largely tolerated in our assay, may in fact be damaging in the endogenous context.

Despite limitations, empirical calibration of our baseline CPOX map showed that it provided new functional evidence for clinical variant classification for 73% of missense variants, which could have immediate clinical value in several scenarios. First, for individuals diagnosed with AHP who meet both clinical and biochemical criteria but carry a CPOX missense variant currently classified as VUS, reclassification towards pathogenicity could enable cascade screening to identify latent HCP in at-risk relatives. Second, when biochemical testing misses the AHP subtype, as can happen for asymptomatic high excretors of ALA and PBG (∼10% of AHP cases)^43^, or is inconclusive, as can happen when it is performed long after the episode, improved pathogenicity classification could enable diagnoses. Third, where biochemical testing has diagnosed an acute attack of porphyria but unable to determine the AHP subtype, improved pathogenicity classification could establish definitive diagnosis. Fourth, for CPOX missense variants identified via direct-to-consumer or newborn genetic screening, the atlas could provide evidence supporting vigilance for HCP symptoms or avoidance of known triggers. Identifying a pathogenic variant can have cascading benefits for potentially-at-risk family members—expanding the number of individuals who benefit from vigilance, preventive measures, or therapy.

Future directions might include assessing quantitative agreement between atlas scores and clinical features such as age of onset or disease severity—which will require establishment of a standardized framework for classifying HCP severity—and ideally stratification by environmental triggers. Additional ‘subfunctional’ assays—measuring variant impacts on protein abundance, specific activity, or mitochondrial localization—could reveal variant mechanisms with different prognosis implications.

We expect that this atlas of CPOX variant impacts will not only yield further insights into sequence-structure-function relationships but also broader therapeutic intervention via more rapid and sensitive genetic diagnosis of HCP and cascade screening of family members.

## 4. METHODS

### Yeast Strains and Plasmids

The *Saccharomyces cerevisiae* strain TSA1076 (*MATα hem13-ts::KanR his3Δ1 leu2Δ0 ura3Δ0*) used to assay activity for *CPOX* variant libraries was kindly provided by Drs. Guihong Tan, Charles Boone, and Brenda Andrews. For yeast expression, wild-type (WT) and mutant *CPOX* open reading frames (ORFs) were subcloned into the Gateway-compatible yeast expression vector pAG415GAL-ccdB (CEN/ARS-based, *GAL1* promoter, and *LEU2* marker) purchased from Addgene. The *CPOX* ORF corresponding to UniprotKB accession P36551 was obtained from Twist Bioscience (South San Francisco, CA, USA) and codon-optimized (via Twist’s default script) for expression in human cells.

Using Gateway LR reactions, both WT and variant *CPOX* ORFs were subcloned into the yeast expression vector (pAG415GAL) and construct sequences were confirmed by Sanger sequencing. Yeast expression vectors (including an empty vector control bearing the ccdB marker under a bacterial promoter) were transformed into bacterial strain NEB DH5ɑ.

### Validation of an environment-dependent and scalable CPOX assays

Yeast ts mutants carrying human *CPOX* WT, variants or empty vector control plasmids were isolated from single colonies and grown to saturation at 30 °C. Each culture was then adjusted to an optical density at 600 nm (OD₆₀₀) of 1.0 and serially diluted in fivefold steps (5^-1^ to 5^-5^). Dilutions (5 µL of each) were spotted onto SC-LEU plates containing 2% galactose as the carbon source to induce *GAL1*-promoter-driven expression and maintain the plasmid. Plates were incubated at either 30 °C or 35 °C for 48 hours. After imaging, results were interpreted by comparing growth of yeast expressing human genes to the corresponding empty vector control (Figure S1A). Two independent cultures corresponding to each distinct plasmid were grown and assayed.

We validated APR measurement in low-throughput liquid growth assays on a Tecan microplate reader by testing CPOX WT, p.Asn272His, and null controls under short-term exposure to moderate mercury concentrations (25–200 µM). Three independent cultures per variant were grown in SC-LEU media with 2% galactose, with or without mercury, at 35 °C overnight to saturation (Figure S1B).

### Generating codon randomized CPOX variant libraries

Mutagenized libraries were generated using a modified version of Precision Oligo-Pool Based Code Alteration (POPCode) approach^44^, as previously described^45^. Mutagenic oligos were designed for each of the 454 codons of *CPOX*, such that each ∼33nt oligo contained a central “NNK” degenerate codon as previously described (see Supplementary Dataset S14 for Primers)^20^.

Three full-length mutagenized construct libraries were generated, with mutagenesis targeted in turn to each of three subregions of the same approximate length. Briefly, from the 454 oligos synthesized (see Supplementary Dataset S14), the mutagenic oligos for each region were combined to produce three regional pools and then phosphorylated. The template plasmid backbone was denatured and the pooled, phosphorylated oligos were annealed together with a primer that hybridized to the 5’ end of the template, adding a single overhang ‘tag’ to allow for preferential amplification of the mutagenized strand by PCR. A fill-in reaction was performed with Phusion MM (NEB), and Taq DNA ligase (NEB) was applied to seal the nicks. Subsequent PCR reactions amplified the mutagenized strand using the tag sequences, and added attB1 and attB2 sites for *en masse* transfer into the entry vector pDONR223 using Gateway BP reactions. Gateway-entry libraries were transformed into NEB DH5-alpha *Escherichia coli* cells (NEB) and selected on LB agar plates containing spectinomycin. After plasmid pool isolation, Gateway-entry clone libraries were transferred to a pAG415GAL expression vector via *en masse* Gateway LR reactions to enable yeast expression. These Gateway-destination libraries were then transformed into NEB DH5-alpha *Escherichia coli* cells (NEB) and selected on LB agar plates containing ampicillin. Next, destination libraries were transformed into the *S. cerevisiae* strain TSA1076 via the EZ Kit Yeast Transformation kit (Zymo Research). To retain high library complexity, plasmids were purified from >500,000 clones at each transfer step and ∼2,000,000 yeast transformants were pooled to form the host library.

### Multiplexed assays for CPOX variant function

High-throughput complementation screening was carried out to assess the functional effects of human CPOX variants in the presence and absence of mercury, as follows: Yeast transformants into the TSA1076 strain were grown at 30°C in synthetic complete medium lacking leucine (SC-LEU; USBiological) with glucose as the carbon source to ensure plasmid retention (pre-selection condition). Plasmid pools were extracted from 10 optical density units (ODUs) of cells (equivalent to the number of yeast cells in 10 mL of a 1.0 OD_600_ culture, typically ∼10⁸ cells) and served as templates for downstream tiling PCR. Two replicates of approximately 4×10⁸ transformants were each washed three times and inoculated into 200 mL SC-LEU medium containing 2% galactose (as the sole carbon source), then incubated at 35°C with or without 96 µM mercury for 48 hours (post-selection conditions). In parallel, the TSA1076 strain was transformed with the WT ORF and grown alongside the variant pool. Plasmids were isolated from 10 ODUs of cells from each culture (two replicates per post-selection condition and WT control) and used as templates for downstream tiling PCR (see Supplementary Dataset S15).

### Sequencing and scoring variant effects

For each plasmid library from both pre-selection and post-selection conditions, short template amplicons (∼150 bp) tiling the *CPOX* ORF (within the context of each regional pool) were amplified using primers containing partial Illumina adaptor sequences (see Supplementary Dataset S15). In a second-round PCR, full-length Illumina sequencing adaptors with sample-specific index tags were added. Paired-end sequencing, which dramatically reduces base-calling errors and enables parts-per-million variant frequencies, was performed for all tiles of pre- and post-selection library. Separate sequencing runs were performed for each assay (with and without mercury) with an Illumina NextSeq 500 via a NextSeq 500/550 High Output Kit v.2, achieving an average sequencing depth of ∼2 million reads paired-end reads per tile. Sequencing reads were demultiplexed with bcl2fastq v.2.17 (Illumina).

Raw read data was processed and variant effects in each assay measured using the previously described TileSeq strategy^19,20^. In brief, variant frequencies in each condition were determined using TileSeq_MutCount (https://github.com/RyogaLi/tileseq_mutcount), which incorporated Bowtie2^46^ for aligning the sequence of each read pair to the reference template. Following alignment, the posterior probability for each divergent base-call was calculated, and only variants exceeding the 0.9 threshold were counted. To account for random error, variants with read counts below 10 or frequencies below the 90th percentile of the WT were filtered out. Functional impact scores were then generated using the tileseqMave pipeline (https://github.com/jweile/tileseqMave). Before calculating enrichment ratios, variant frequencies in both the pre- and post-selection libraries were adjusted by subtracting the corresponding WT frequencies. Each variant’s score was derived by comparing its relative enrichment to the median ratios observed for synonymous and nonsense variants. For both assays, the scores were rescaled for each region separately such that, post-rescaling, the medians of nonsense and synonymous variants were 0 and 1, respectively. Measurement uncertainty was regularized using the method of Baldi and Long^47^ and propagated through calculations via bootstrapping, as previously described^7^.

Variants were initially classified as intolerant to variation if the baseline score plus its standard error was below 0.5 in either the baseline or mercury map. This threshold corresponded to the local minimum between the two modes in the bimodal distribution of missense variant scores.

We calculated a delta score (mercury – baseline) for each variant using the unscaled map scores to account for uniform mercury-dependent inhibition. Standard errors were computed using error propagation, and delta scores were considered significant if their 95% confidence intervals excluded zero. A two-sided Wilcoxon signed-rank test was used to test the alternative hypothesis that mercury scores are significantly different from baseline scores. Nonsense and synonymous variants served as control sets to derive a null distribution for delta scores. To isolate missense variants with altered substrate preference, we first identified those with significant delta scores, defined as variants with 95% confidence intervals that excluded the null distribution median. We then filtered this set to include only variants considered as tolerated in the baseline map.

### Relating map scores to population allele frequencies

Non-overlapping sets of *CPOX* variant allele frequencies were obtained from population databases, including gnomAD v4.1.0 (https://gnomad.broadinstitute.org/) and the UK Biobank (OQFE whole-exome VCFs; application ID: 51135), which includes sequencing data from ∼450,000 participants. We then calculated odds ratios to assess allele depletion for variants classified as either damaging or neutral in our baseline map.

### Interpreting hyper-complementation scores

To determine how hyper-complementing variants (those with greater-than–wild-type functional scores) in our baseline map are best interpreted in a human context, we performed phylogenetic analysis as previously described^19,20,48^. Three models relating variant scores to evolutionary amino acid preference were evaluated: proportional to the experimental score (hyper-complementing is advantageous), capped at the wild-type score (hyper-complementing is neutral), or set to the reciprocal of scores exceeding wild-type (hyper-complementing is deleterious). Each model was tested on 33 Ensembl homologs with ≥85% sequence identity to the human protein (Supplementary Dataset S12) using *phydms* (https://github.com/jbloomlab/phydms), and fit to the CPOX phylogeny was assessed via the Akaike information criterion to identify the best-fitting model.

### Normalization and transformation of AlphaMissense scores

Scores from the baseline variant effect map were placed on a common scale with those of the computational predictor AlphaMissense^38^ using *s*_tranformed_ = 1 − (*s*_original_ − min(*s*))/(max(*s*) − min(*s*)), such that, after transformation, 0 represents null-like variants, while 1 represents neutral variants.

### Solvent accessibility and thermostability calculations

We used the FreeSASA program (https://freesasa.github.io/) to calculate the relative solvent exposure of residue positions. After examining the distribution of relative solvent exposure values, we established thresholds corresponding to the high and low peaks. Residues with surface area values exceeding 50% were considered exposed, while those below 25% were classified as buried.

To determine whether our atlas of functional scores reflect changes in steady-state protein levels (i.e., stability) rather than specific activity, we calculated protein thermostability (ΔΔG) values using DDGun3D version 0.0.2 (https://github.com/biofold/ddgun), as previously described^49^. The PDB entry 2AEX for CPOX satisfied the following criteria: an X-ray determined structure with resolution 1.58 Å or better, homodimer structure, and no missing or non-standard residues^50^. We defined stabilizing amino acid substitutions as those for which ΔΔG ≥ −0.1, and a destabilizing substitution for ΔΔG < −0.1.

### MD and protein-ligand docking simulations

#### CPOX structures and models

The human CPOX crystal structure was obtained from the Protein Data Bank^51^ (PDB ID: 2AEX^21^), a homodimer complexed with citric acid (CIT). The AlphaFold model was generated by using two identical sequences from the UniProt database^52^ (UniProt ID: P36551) including the ligand CIT in the complex. AlphaFold (AF) models were generated using the AlphaFold3 web server (https://alphafoldserver.com) and the top model was selected for simulations^53^. Variants p.Asn272His, p.Gly188Gln, p.Val135Ala and p.Leu155Trp were introduced into both the crystal and AF structures using the PyMOL Mutagenesis tool (https://www.pymol.org/).

#### Determination of hinge sites

Hinge residues were identified from the Gaussian Network Model (GNM)^3030^ normal modes by analyzing the normalized displacement profiles of residues along each mode. For global modes, residues located at sign crossovers of the eigenvector components (transitions between positive and negative fluctuations) within a narrow band around zero are defined as hinge sites, as they undergo minimal translations while mediating the opposite movements of domains/chains on both sides. These positions often correspond to functionally important mechanical control points in protein motions.

#### MD simulations on apo WT and variants

The solvated systems and inputs for NAMD simulations were prepared using the CHARMM-GUI Solution Builder module (with default parameters/procedure)^54,55^. A rectangular water box of 10 Å edge distance was constructed around each system. Physiological conditions (0.15 M KCl, pH 7.4) were maintained. The CHARMM36m force field was used^56^. NPT ensemble was used for the production phase (P= 1 atm; T = 303.15 K). Details of multiple independent runs are given in Table S2.

#### Analysis of CPOX dynamics and secondary structural changes

MD snapshots were recorded every 100 ps during each run. ProDy API was used to perform principal component analysis (PCA) after aligning WT and variant snapshots onto the starting structure (initial AF model or crystal structure)^57^. Additionally, we calculated the root mean square deviations (RMSDs) of snapshots with respect to the reference structure and the residue mean square fluctuations (MSFs) for WT and each variant. We also analyzed the changes in secondary structural elements using the VMD secondary structure evaluation module (see Supplementary Dataset S16)

#### Analysis of active site volumes and gates

To identify MD snapshots (extracted by the MDAnalysis tool)^58^ which could potentially accommodate COPRO and/or URO binding, we employed two criteria: First, the “volumes” criterion was based on the estimated size of the active site pocket volume in each snapshot. Using the FPocket algorithm, we detected all pockets and then identified the active site pockets—one on each chain—by selecting those with the highest number of substrate-interacting residues. Since no structure has been resolved with the substrate, we used the residues interacting with CIT (PDB ID: 2AEX) as our reference. The corresponding active site pocket volumes were then extracted for further analysis.

The second “gates” criterion was based on the estimated sizes of each of three potential entrances we identified to the active site (described as Gates 1, 2, and 3). Whether each gate is open in each snapshot was determined via measurements of distance between specific residue pairs (Figures 5 and S10; see Table S4 for thresholds and details). Finally, to identify MD snapshots capable of successful docking, we combined these criteria to require that at least one gate be open and that the pocket volume was sufficiently large (> 2000 Å^3^). Among the snapshots with at least one open gate, those with the highest pocket volumes (top 2 or 3) were selected for substrate docking.

#### Substrate Docking

Autodock Vina^59^ with default settings was used for COPRO and URO dockings onto the active site. In each case we used at least two snapshots, each having at least one open active site pocket. The docking box was set to encompass the interface and both active sites of dimeric CPOX. Each docking simulation consisted of five independent runs of 20 poses (100 poses in total). The exhaustiveness of each run was set to 50 (from default 8) for broad sampling of potential binding conformations for each ligand.

#### MD simulations on docked poses

To observe the persistence of bound ligands and the binding affinity of the protein-ligand poses, we modeled each complex in Amber 24^60–62^ (forcefield: protein ff19SB^63^, ligands GAFF2^64^ with AM1-BCC charges^65^) using octahedral periodic boundary conditions (10 Å buffer around the protein, TIP3P water model^66^, neutralized^66,67^).Each system was then minimized (four stages), heated to 300 K, and equilibrated twice under NPT by gradually reducing ligand restraints. Production MD runs (Table S2) were performed with 2 fs timesteps under NPT (Langevin dynamics with γ=1 ps⁻¹, non-bonded interaction cutoff=9Å) conditions^68,69^.

#### Binding affinity estimation using PRODIGY-LIG

To estimate binding affinity based on MD simulations, we ran PRODIGY-LIG^33,70^ on MD frames (100 ps apart), scoring for each ligand bound to either chain A or B (depending on the docking score, see Table S4). For each system, the ligand-bound chain and binding pose used for subsequent analysis were selected from the AutoDock Vina docking results based on the consensus between the most favorable docking score and the highest active-site pocket occupancy. The selected docking-defined chain was then used consistently for PRODIGY-LIG analysis of the corresponding ligand-bound holo MD trajectory frames, generating a single time-dependent binding free energy profile, ΔG(t), for subsequent analysis.

### A reference set of clinically annotated variants

To evaluate the ability of variant effect maps to identify pathogenic variants, we compiled a “positive” set of twenty-one disease-associated variants (sixteen HCP-causing and five HP-causing) curated by expert centers in Europe and the United States and classified via Labcorp Genetics’ Sherloc v.6.0 framework^71^ and ClinVar^13^. For all ClinVar-submitted variants, review criteria had been provided and there were no conflicting interpretations.

### Calibrating Log-Likelihood Ratios to ACMG/AMP Evidence Strengths

To assess the strength of evidence supporting pathogenicity for each variant, we calculated the log-likelihood ratio of pathogenicity (LLRp) corresponding to each functional impact score from the baseline map. Briefly, the scores from positive and negative reference variants were separately used to estimate probability density functions using kernel density estimation, applying a Gaussian kernel with bandwidth selected via biased cross-validation. For each variant, the LLRp was calculated as the log-ratio of the densities from pathogenic and benign distributions, respectively, with regularization to a uniform (LLRp = 0) background to mitigate overfitting in regions with sparse data. These LLRp values were then classified using ACMG/AMP guidelines using the method of Tavtigian et al. ^42^, as later modified^20^.

## Data code and availability

Functional impact scores from both maps, as well as baseline map LLRp values with confidence intervals and ACMG-compatible evidence strengths, are available in the Supplementary Dataset. Scores can also be accessed on MaveDB^72^ for both the baseline map (accession number: urn:mavedb:00001282-a-1) and the mercury map (accession number: urn:mavedb:00001282-a-2). NHGRI’s [Impact of Genomic Variation on Function (IGVF) Consortium standards documents](https://data.igvf.org/search/?type=AnalysisSet&status!=deleted&lab.title=Frederick+Roth%2C+UToronto) include links to protocols.io^73^, detailed quality control metrics and raw data. Custom scripts for all downstream analyses are publicly available on GitHub: https://github.com/HaotianFrankZhang/CPOX.git

## Web resources

TileSeq_MutCount, https://github.com/RyogaLi/tileseq_mutcount

TileSeqMave, https://github.com/jweile/tileseqMave

Phydms, https://github.com/jbloomlab/phydms

## Acknowledgements

We are grateful to J. Reuter for facilitating collaboration with Labcorp Genetics (formerly Invitae Corp.). We thank G. Lum and A. Nasrabad for computational assistance with molecular dynamics simulations. We also thank the participants of the UK Biobank, as well as the individuals who envisioned, developed, and continue to support these valuable resources. Access to the UK Biobank was provided under a material transfer agreement with Sinai Health.

## Funding

University of Toronto Precision Medicine Initiative (PRiME) Fellowship PRMF2020 (to W.vL.). National Human Genome Research Institute of the National Institutes of Health Center of Excellence in Genomic Science Initiative RM1HG010461 (to F.P.R.). NHGRI Impact of Genomic Variation on Function Initiative UM1HG011989 (to F.P.R.). Canadian Institutes of Health Research Foundation Grant FDN159926 (to F.P.R.). National Institute of General Medical Sciences R01 GM139297 (to I.B. and P.D.).

## Author contributions

Conceptualization: J.TF., W.vL., and F.P.R. Methodology: W.vL., H.Z., P.D., and F.P.R. Investigation: W.vL., H.Z., V.S., M.P., A.R., A.A., M.G., R.J.D, B.W., C.S., J.TF., A.W., and F.P.R. Analysis and visualization: W.vL., H.Z., P.D., and F.P.R. Funding acquisition: W.vL and F.P.R. Supervision: F.P.R. and I.B. Writing (original draft): W.vL., H.Z., P.D., and F.P.R. Writing (review & editing): W.vL., H.Z., V.S., M.J.C., R.J.D., C.S., J.TF., A.W., I.B., P.D., and F.P.R.

## Competing interests

F.P.R. is an investor in Ranomics, Inc., and is an investor in and advisor for SeqWell, Inc. and Constantiam Biosciences, Inc.

## Notes

### Competing Interest Statement

The authors have declared no competing interest.

## REFERENCES

1. Weile, J. & Roth, F. P. Multiplexed assays of variant effects contribute to a growing genotype-phenotype atlas. Hum. Genet. 137, 665–678 (2018).

2. Pérez-Palma, E., Gramm, M., Nürnberg, P., May, P. & Lal, D. Simple ClinVar: an interactive web server to explore and retrieve gene and disease variants aggregated in ClinVar database. Nucleic Acids Res. 47, W99–W105 (2019).

3. Richards, S. et al. Standards and guidelines for the interpretation of sequence variants: a joint consensus recommendation of the American College of Medical Genetics and Genomics and the Association for Molecular Pathology. Genet. Med. 17, 405–424 (2015).

4. Fayer, S. et al. Closing the gap: Systematic integration of multiplexed functional data resolves variants of uncertain significance in BRCA1, TP53, and PTEN. Am. J. Hum. Genet. 108, 2248–2258 (2021).

5. Floyd, B. J. et al. Proactive variant effect mapping aids diagnosis in pediatric cardiac arrest. Circ. Genom. Precis. Med. 16, e003792 (2023).

6. Tejura, M., et al. A scalable approach to resolving variants of uncertain significance. bioRxivorg (2026) doi:10.64898/2026.02.14.705848.

7. Weile, J. et al. Shifting landscapes of human MTHFR missense-variant effects. Am. J. Hum. Genet. 108, 1283–1300 (2021).

8. Flynn, J. M. et al. Comprehensive fitness maps of Hsp90 show widespread environmental dependence. Elife 9, (2020).

9. Thompson, S., Zhang, Y., Ingle, C., Reynolds, K. A. & Kortemme, T. Altered expression of a quality control protease in E. coli reshapes the in vivo mutational landscape of a model enzyme. Elife 9, (2020).

10. Martin-Rufino, J. D. et al. Massively parallel base editing to map variant effects in human hematopoiesis. Cell 186, 2456–2474.e24 (2023).

11. Phillips, J. D. Heme biosynthesis and the porphyrias. Mol. Genet. Metab. 128, 164–177 (2019).

12. Wang, B., Bonkovsky, H. L., Lim, J. K. & Balwani, M. AGA clinical practice update on diagnosis and management of acute hepatic porphyrias: Expert review. Gastroenterology 164, 484–491 (2023).

13. Landrum, M. J. et al. ClinVar: public archive of relationships among sequence variation and human phenotype. Nucleic Acids Res. 42, D980–5 (2014).

14. Karczewski, K. J. et al. The mutational constraint spectrum quantified from variation in 141,456 humans. Nature 581, 434–443 (2020).

15. Woods, J. S. et al. Modification of neurobehavioral effects of mercury by a genetic polymorphism of coproporphyrinogen oxidase in children. Neurotoxicol. Teratol. 34, 513–521 (2012).

16. Heyer, N. J., Bittner, A. C., Jr, Echeverria, D. & Woods, J. S. A cascade analysis of the interaction of mercury and coproporphyrinogen oxidase (CPOX) polymorphism on the heme biosynthetic pathway and porphyrin production. Toxicol. Lett. 161, 159–166 (2006).

17. Woods, J. S. et al. The association between genetic polymorphisms of coproporphyrinogen oxidase and an atypical porphyrinogenic response to mercury exposure in humans. Toxicol. Appl. Pharmacol. 206, 113–120 (2005).

18. Kachroo, A. H. et al. Systematic bacterialization of yeast genes identifies a near-universally swappable pathway. Elife 6, (2017).

19. Weile, J. et al. A framework for exhaustively mapping functional missense variants. Mol. Syst. Biol. 13, 957 (2017).

20. van Loggerenberg, W. et al. Systematically testing human HMBS missense variants to reveal mechanism and pathogenic variation. Am. J. Hum. Genet. 110, 1769–1786 (2023).

21. Lee, D.-S. et al. Structural basis of hereditary coproporphyria. Proc. Natl. Acad. Sci. U. S. A. 102, 14232–14237 (2005).

22. Susa, S. et al. The long, but not the short, presequence of human coproporphyrinogen oxidase is essential for its import and sorting to mitochondria. Tohoku J. Exp. Med. 200, 39–45 (2003).

23. Singh, A. & Xu, Y.-J. Heme deficiency sensitizes yeast cells to oxidative stress induced by hydroxyurea. J. Biol. Chem. 292, 9088–9103 (2017).

24. Grandchamp, B., Phung, N. & Nordmann, Y. The mitochondrial localization of coproporphyrinogen III oxidase. Biochem. J. 176, 97–102 (1978).

25. Schmitt, C. et al. Mutations in human CPO gene predict clinical expression of either hepatic hereditary coproporphyria or erythropoietic harderoporphyria. Hum. Mol. Genet. 14, 3089–3098 (2005).

26. Moghe, A., Ramanujam, V. M. S., Phillips, J. D., Desnick, R. J. & Anderson, K. E. Harderoporphyria: Case of lifelong photosensitivity associated with compound heterozygous coproporphyrinogen oxidase (CPOX) mutations. Mol. Genet. Metab. Rep. 19, 100457 (2019).

27. Hasanoglu, A. et al. Harderoporphyria due to homozygosity for coproporphyrinogen oxidase missense mutation H327R. J. Inherit. Metab. Dis. 34, 225–231 (2011).

28. Li, T. & Woods, J. S. Cloning, expression, and biochemical properties of CPOX4, a genetic variant of coproporphyrinogen oxidase that affects susceptibility to mercury toxicity in humans. Toxicol. Sci. 109, 228–236 (2009).

29. To-Figueras, J. et al. Biochemical and genetic characterization of four cases of hereditary coproporphyria in Spain. Mol. Genet. Metab. 85, 160–163 (2005).

30. Zhang, H., Gur, M. & Bahar, I. Global hinge sites of proteins as target sites for drug binding. Proc. Natl. Acad. Sci. U. S. A. 121, e2414333121 (2024).

31. Borrero Corte, M. J., et al. Molecular analysis of 19 Spanish patients with mixed porphyrias. Eur. J. Med. Genet. 62, 103589 (2019).

32. Phillips, J. D. et al. Crystal structure of the oxygen-dependant coproporphyrinogen oxidase (Hem13p) of Saccharomyces cerevisiae. J. Biol. Chem. 279, 38960–38968 (2004).

33. Vangone, A. et al. Large-scale prediction of binding affinity in protein-small ligand complexes: the PRODIGY-LIG web server. Bioinformatics 35, 1585–1587 (2019).

34. Pejaver, V. et al. Calibration of computational tools for missense variant pathogenicity classification and ClinGen recommendations for PP3/BP4 criteria. Am. J. Hum. Genet. 109, 2163–2177 (2022).

35. Bergquist, T. et al. Calibration of additional computational tools expands ClinGen recommendation options for variant classification with PP3/BP4 criteria. bioRxivorg 2024.09.17.611902 (2024).

36. Livesey, B. J. & Marsh, J. A. Variant effect predictor correlation with functional assays is reflective of clinical classification performance. Genome Biol. 26, 104 (2025).

37. Livesey, B. J. & Marsh, J. A. Updated benchmarking of variant effect predictors using deep mutational scanning. Mol. Syst. Biol. 19, e11474 (2023).

38. Cheng, J. et al. Accurate proteome-wide missense variant effect prediction with AlphaMissense. Science 381, eadg7492 (2023).

39. Wu, Y., Li, R., Sun, S., Weile, J. & Roth, F. P. Improved pathogenicity prediction for rare human missense variants. Am. J. Hum. Genet. 108, 1891–1906 (2021).

40. Brandes, N., Goldman, G., Wang, C. H., Ye, C. J. & Ntranos, V. Genome-wide prediction of disease variant effects with a deep protein language model. Nat. Genet. (2023) doi:10.1038/s41588-023-01465-0.

41. Tavtigian, S. V., Harrison, S. M., Boucher, K. M. & Biesecker, L. G. Fitting a naturally scaled point system to the ACMG/AMP variant classification guidelines. Hum. Mutat. 41, 1734–1737 (2020).

42. Tavtigian, S. V. et al. Modeling the ACMG/AMP variant classification guidelines as a Bayesian classification framework. Genet. Med. 20, 1054–1060 (2018).

43. Balwani, M. et al. Acute hepatic porphyrias: Recommendations for evaluation and long-term management. Hepatology 66, 1314–1322 (2017).

44. Axakova, A. et al. Landscapes of missense variant impact for human superoxide dismutase 1. Genetics (2025).

45. Zimmerman, D., et al. Comprehensively testing the function of missense variation in the STK11 tumour suppressor. bioRxivorg (2025) doi:10.1101/2025.07.14.664734.

46. Langmead, B. & Salzberg, S. L. Fast gapped-read alignment with Bowtie 2. Nat. Methods 9, 357–359 (2012).

47. Baldi, P. & Long, A. D. A Bayesian framework for the analysis of microarray expression data: regularized t -test and statistical inferences of gene changes. Bioinformatics 17, 509–519 (2001).

48. Hilton, S. K., Doud, M. B. & Bloom, J. D. phydms: software for phylogenetic analyses informed by deep mutational scanning. PeerJ 5, e3657 (2017).

49. Montanucci, L., Capriotti, E., Frank, Y., Ben-Tal, N. & Fariselli, P. DDGun: an untrained method for the prediction of protein stability changes upon single and multiple point variations. BMC Bioinformatics 20, 335 (2019).

50. Lee, D. S. et al. The 1.58A crystal structure of human coproporphyrinogen oxidase reveals the structural basis of hereditary coproporphyria. Preprint at 10.2210/pdb2aex/pdb (2005).

51. Burley, S. K. et al. RCSB Protein Data Bank: powerful new tools for exploring 3D structures of biological macromolecules for basic and applied research and education in fundamental biology, biomedicine, biotechnology, bioengineering and energy sciences. Nucleic Acids Res. 49, D437–D451 (2021).

52. UniProt Consortium. UniProt: The universal protein knowledgebase in 2023. Nucleic Acids Res. 51, D523–D531 (2023).

53. Abramson, J. et al. Addendum: Accurate structure prediction of biomolecular interactions with AlphaFold 3. Nature 636, E4 (2024).

54. Jo, S., Kim, T., Iyer, V. G. & Im, W. CHARMM-GUI: a web-based graphical user interface for CHARMM. J. Comput. Chem. 29, 1859–1865 (2008).

55. Phillips, J. C. et al. Scalable molecular dynamics on CPU and GPU architectures with NAMD. J. Chem. Phys. 153, 044130 (2020).

56. Lee, J. et al. CHARMM-GUI input generator for NAMD, GROMACS, AMBER, OpenMM, and CHARMM/OpenMM simulations using the CHARMM36 additive force field. J. Chem. Theory Comput. 12, 405–413 (2016).

57. Bakan, A. et al. Evol and ProDy for bridging protein sequence evolution and structural dynamics. Bioinformatics 30, 2681–2683 (2014).

58. Michaud-Agrawal, N., Denning, E. J., Woolf, T. B. & Beckstein, O. MDAnalysis: a toolkit for the analysis of molecular dynamics simulations. J. Comput. Chem. 32, 2319–2327 (2011).

59. Trott, O. & Olson, A. J. AutoDock Vina: improving the speed and accuracy of docking with a new scoring function, efficient optimization, and multithreading. J. Comput. Chem. 31, 455–461 (2010).

60. Case, D. A. et al. Amber 2025. (2025).

61. Case, D. A. et al. AmberTools. J. Chem. Inf. Model. 63, 6183–6191 (2023).

62. Case, D. A. et al. Recent developments in Amber biomolecular simulations. J. Chem. Inf. Model. 65, 7835–7843 (2025).

63. Tian, C. et al. Ff19SB: Amino-acid-specific protein backbone parameters trained against quantum mechanics energy surfaces in solution. J. Chem. Theory Comput. 16, 528–552 (2020).

64. Development and testing of a general amber force field after AM1-BCC, please cite this paper: Fast, efficient generation of high-quality atomic charges. in AM1-BCC model: II. Parameterization and validation.

65. Jakalian, A., Jack, D. B. & Bayly, C. I. Fast, efficient generation of high-quality atomic charges. AM1-BCC model: II. Parameterization and validation. J. Comput. Chem. 23, 1623–1641 (2002).

66. Jorgensen, W. L., Chandrasekhar, J., Madura, J. D., Impey, R. W. & Klein, M. L. Comparison of simple potential functions for simulating liquid water. J. Chem. Phys. 79, 926–935 (1983).

67. Determination of Alkali and Halide Monovalent Ion Parameters for TIP3P Water Langevin Dynamics Is Good.

68. Leimkuhler, B. & Matthews, C. Robust and efficient configurational molecular sampling via Langevin dynamics. J. Chem. Phys. 138, 174102 (2013).

69. Loncharich, R. J., Brooks, B. R. & Pastor, R. W. Langevin dynamics of peptides: the frictional dependence of isomerization rates of N-acetylalanyl-N’-methylamide. Biopolymers 32, 523–535 (1992).

70. Kurkcuoglu, Z. et al. Performance of HADDOCK and a simple contact-based protein-ligand binding affinity predictor in the D3R Grand Challenge 2. J. Comput. Aided Mol. Des. 32, 175–185 (2018).

71. Nykamp, K. et al. Correction: Sherloc: a comprehensive refinement of the ACMG-AMP variant classification criteria. Genet. Med. 22, 240 (2020).

72. Esposito, D. et al. MaveDB: an open-source platform to distribute and interpret data from multiplexed assays of variant effect. Genome Biol. 20, 223 (2019).

73. van Loggerenberg, W., Zimmerman, D., Axakova, A. & de Souza, A. Variant Effect mapping protocol collection v1. (2025) doi:10.17504/protocols.io.5qpvo1n37g4o/v1.

