## Supplementary Notes and Figures for "Environment-dependent landscapes of coding variant impacts on coproporphyrinogen oxidase"

#### 1. Hyper-complementing CPOX variants are likely detrimental in humans

Human missense variants showing hyper-complementation in a yeast-based assay can for some genes and assays tend to be detrimental in humans, as described previously for *UBE2I*[1,2] and more recently for HMBS, the sixth enzyme in the heme biosynthetic pathway. This was shown via tests of whether or not variants are more likely or less likely to be observed in the orthologous metazoan genes. More specifically, the set of apparently hyper-complementing variants is tested for goodness of fit for each of three phylogenetic models[3,4]: (1) hyper-complementing variants confer a fitness advantage in humans and related species; (2) hyper-complementing variants are functionally equivalent to wild-type (WT); or (3) hyper-complementing variants are detrimental in humans with a fitness effect that follows the reciprocal of the yeast-assay-derived functional impact score. When applied to our CPOX scores (the baseline map), this analysis showed that hyper-complementing variants were less likely than other variants to appear in related species, supporting the third model (Table S1) in which apparently hypercomplementing variants have an *in vivo* fitness disadvantage. We also evaluated this question using a top-performing computational variant effect predictor (AlphaMissense[5]): After first transforming AlphaMissense scores to the same scale as our baseline map (see Material and Methods), we found that AlphaMissense scored hyper-complementing variants as being more detrimental than those with WT-like activity (Figure S4). Together, these findings support the conclusion that observation of hyper-complementation in our assay should be taken as evidence towards detrimental effects in humans.

#### 2. Active-site loop conformational variability: unstructured vs. helical states

Human CPOX is active as a homodimer (Figure S9), with each monomer adopting a flat seven-stranded β-sheet flanked by α-helices. This unusual flatness has been proposed to result from the high glycine content of β2–4, 6, and 7[6], with the active site positioned between the β-sheet and helices α9–α11, where His258 (β5) has a critical role in catalysis and Arg262 (β5) and Ser244 (β4) contribute to substrate recognition.

The largest conformational variation between crystal structures of human CPOX (PDB ID: 2AEX)[6] and the yeast ortholog Hem13 is the active site loop which lines the upper part of the active site cavity: it is largely unstructured in the human crystal but forms a stable helix (α10) in the yeast ortholog, a feature also captured by the AlphaFold3 (AF) model (see Figure 3). Despite these differences, the residues comprising the active site loop are largely conserved in CPOX[7]. Yeast Hem13 has been solved both in the open state (PDB: 1TKL; Figure S1E) and the closed state (PDB: 1TLB; Figure S1F), revealing lid (α4) movements of ~10 Å (A82-F279 distance) relative to α10 that enable substrate access[8]. To comprehensively assess both loop conformations, we performed MD simulations starting from the human crystal structure with the unstructured loop. In the following sections, we present functional dynamics and active site properties for crystal simulations in comparison to AF-based simulations with the helical loop (see main text). We also note that the loop retains its initial conformation—whether helical or unstructured—throughout simulations, without transitioning between the two states.

#### 2A. Substrate accessibility decreases when the active site loop is unstructured

We observed the active site pocket volumes to be markedly reduced in the presence of the unstructured loop (Figure S10). Median volumes for WT and all variants range from 655–742 Å³ in crystal simulations, compared to 1609–1936 Å³ in AF simulations with the helical loop (Figure S10B). This stark difference is further illustrated by the contrast between the initial pocket (573 Å³) and a rarely sampled, more open pocket during crystal simulations (2310 Å³; Figure S10A). Thus, the large COPRO and URO substrates (Figure S10C) could not be accommodated within the active site in most snapshots featuring the unstructured loop.

Significant changes were also observed in the active site gates when the loop was unstructured (Figure S10D, E). While gate 1 remained mostly closed, gate 2, together with gate 3, appeared to serve as the primary route for ligand entry. This is in contrast to AF simulations, where coordinated opening of gates 1 and 2, enabled by the structured lid and helical loop, led to more accessible and spacious active sites. Collectively, these results indicate that the unstructured loop restricts substrate access and reduces the likelihood of sampling open, docking-competent conformations.

**2B. Substrate preferences via docking in the presence of unstructured loop**

Docking results for the snapshots from the yeast CPOX ortholog Hem13 structure (Table S3) reveal notable differences in substrate preferences (COPRO vs URO) between WT and variants. WT Hem13 with its unstructured loop shows a clear preference for COPRO (−9.6 kcal/mol) over URO (−7.9 kcal/mol; Δ = −1.7 kcal/mol), supporting its native substrate specificity. However, low docking occupancies for both COPRO (10%) and URO (5%) indicate that the unstructured loop creates a generally unfavorable binding environment. In contrast, variants shift preference toward URO, with higher docking occupancies and comparable or stronger binding affinities, reflecting reduced specificity for COPRO and suggesting that these mutations promote binding of non-native substrates. The overall picture that emerges from substrate dockings to Hem13 with the unstructured active site loop is consistent with the preferences observed for the helical loop in AF snapshots from human CPOX (see main text). These trends suggest that mutations disrupt the balance for selective binding, leading to increased susceptibility to undesired products.

**2C. Comparison of CPOX dynamics with helical vs unstructured loop**

Mean-square fluctuation (MSF) analysis (Figure S11A) shows that both the lid (α4) and active-site loop exhibit higher fluctuations when the active site loop is structured. Thus, there is higher mobility in the presence of α10 helix (AF model), compared to the unstructured loop (crystal structure). Furthermore, RMSD distributions (Figure S11B,C) reveal consistently higher values in AF simulations for both WT and all variants, relative to crystal simulations. These results explain the greater conformational variability and increased tendency to adopt open active site conformations in the AF model.

We determined the dynamic domains using the SPECTRUS[9] server (Figures S11D,E), which decomposes the structure into quasi-rigid domains based on GNM. In the crystal structure, the active site loop and lid domain formed a single dynamic unit, suggesting that they move in a coordinated manner. This coupled motion restricted ligand entrance by keeping gate 1 in the closed conformation. In contrast, the AF model classified these regions as separate dynamic domains with independent movement. This decoupling appeared to facilitate gate 1 opening and substrate entrance to the active site.

**2D. MD simulations on CPOX-substrate complexes**

To assess the persistence of docked substrates in the active site, 200 ns molecular dynamics simulations were initiated from the top-ranked poses, with two independent runs for the crystal structure and three for the AF model (Table S2). Across all simulations, the substrates remained stably confined within the binding pocket throughout the 200 ns trajectories.

Binding affinities of the protein-ligand complexes were estimated using PRODIGY-LIG[10] for 2000 frames sampled along each trajectory, with the best score selected between the two chains for each snapshot. WT Hem13 showed slightly stronger binding to COPRO than to URO, whereas this trend was reversed for the p.Asn272His variant, which favored URO (Figure S12). In AF-based simulations, the variants consistently shifted substrate preference from COPRO toward URO, in agreement with our initial hypothesis.

**2E. Evaluating the effects of variants on putative mercury-binding sites**

To investigate whether variants alter the accessibility of potential mercury-binding sites in CPOX, we evaluated the solvent-accessible surface area (SASA) of all cysteine residues across WT and mutant (p.Leu155Trp, p.Gly188Gln, and p.Val135Ala and p.Asn272His) apo MD simulations. SASA values were calculated using the PyMOL *get_area* function with solvent-accessible surface settings (dot_solvent = 1, water probe radius = 1.4 Å, and dot density = 3). All ten cysteine residues were analyzed separately in both chains, and SASA values were reported as absolute surface areas (Å²) (Figure S13).

Across all cysteine residues, SASA distributions were broadly similar between WT and variants, with only minor mutation-dependent differences observed (Figure S13). These findings suggest that the variants examined here have limited effects on the solvent accessibility of candidate mercury-reactive cysteine residues.

Among these residues, Cys319 is of particular interest because mercury binding at this site has been proposed to allosterically alter substrate affinity[11]. Consistent with its location adjacent to the active site and its contribution to Gate 2 formation (Figure 5A and Figure S10D), Cys319 remained relatively solvent-accessible throughout the simulations (Figure S13C). However, the variants produced only modest changes in Cys319 SASA, consistent with the broader pattern observed across all cysteine residues. Notably, our baseline map indicates that Cys319 is highly functionally constrained, with substitutions at this position (in either baseline or mercury map) predicted to be strongly deleterious. Together, these findings support an important structural and functional role for Cys319 in CPOX while suggesting that the variants examined here are unlikely to act primarily through substantial changes in Cys319 exposure/accessibility to mercury.

Taken altogether, our results suggest that, while both mercury and many variants can cause CPOX dysfunction, these impacts are not necessarily mechanistically coupled. Candidate mercury-binding cysteines remained largely solvent-accessible across simulations of mercury-sensitive variants, suggesting that these variants do not substantially alter mercury-binding-site exposure. Instead, variants appeared to exert their effects through changes in substrate binding, gate dynamics, and active-site organization. This interpretation is consistent with previous biochemical characterization of p.Asn272His, which showed reduced substrate affinity and diminished basal enzymatic activity, while $Hg^{2+}$ produced additional inhibition of enzyme function [12]. Accordingly, our simulations focused on apo and substrate-bound states to examine how variants reshape intrinsic protein dynamics.

### Supplementary Figures

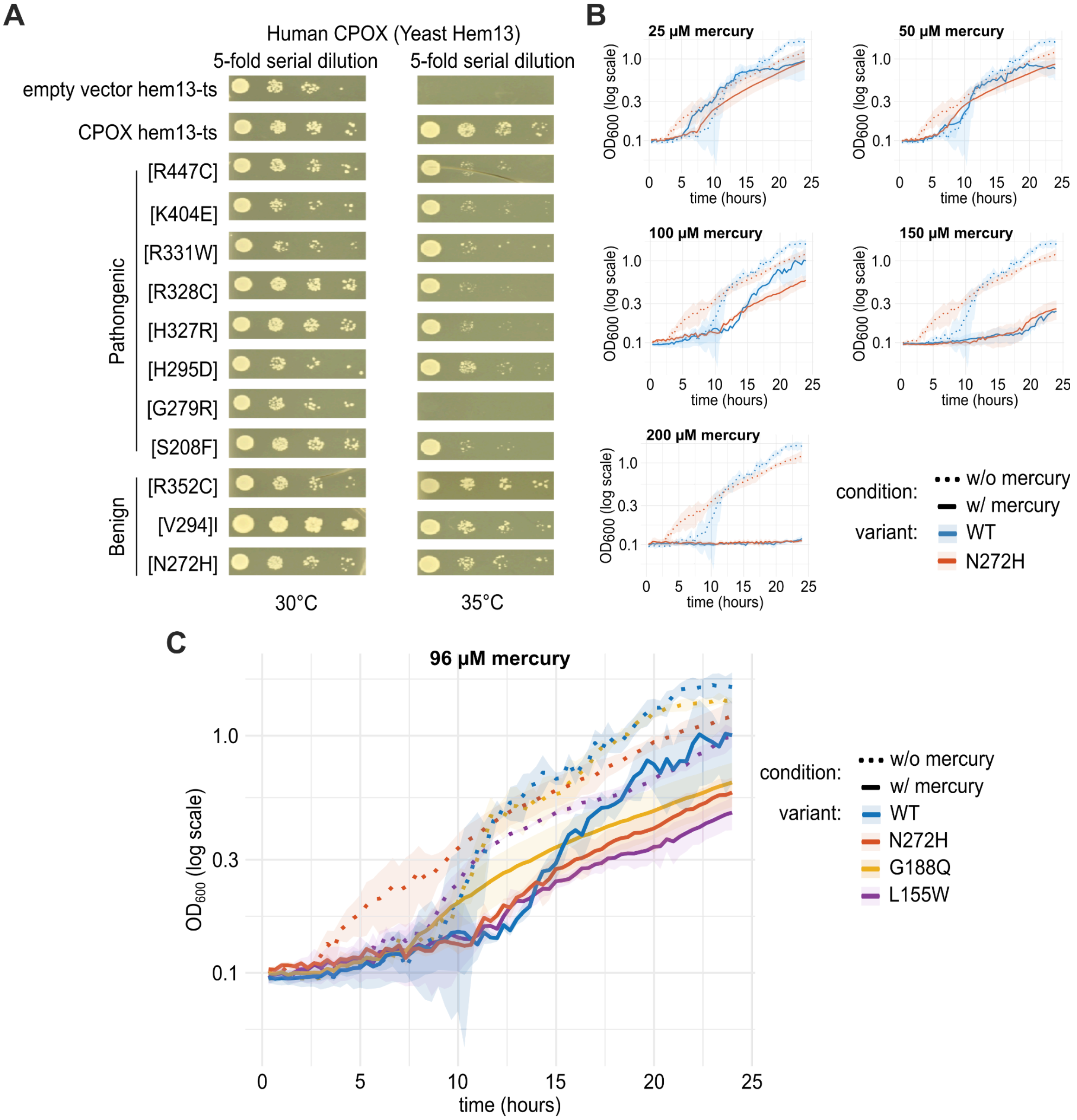


**Figure S1.** Functional complementation assays assessing whether human CPOX variants rescue growth of a yeast strain bearing a temperature-sensitive (ts) mutation in the essential gene *hem13*, either in the presence or absence of mercury. (A) Spotting assay validation of functional complementation for ClinVar-reported pathogenic and benign variants, along with WT and empty-vector controls. Five-fold serial dilutions were spotted onto plates and grown for 48 hours at permissive (30°C) or non-permissive (35°C) temperatures. (B) Liquid growth assay confirmation. Continuous optical density measurements (λ = 600 nm) were obtained for the yeast *hem13* ts strain expressing WT human CPOX or p.Asn272His across five mercury concentrations (25, 50, 96, 150, and 200 µM) at nonpermissive temperature. (C) Validation of a severe functional defect associated with a mercury-induced atypical porphyrinogenic response for variants p.Asn272His, p.Leu155Trp and p.Gly188Gln.

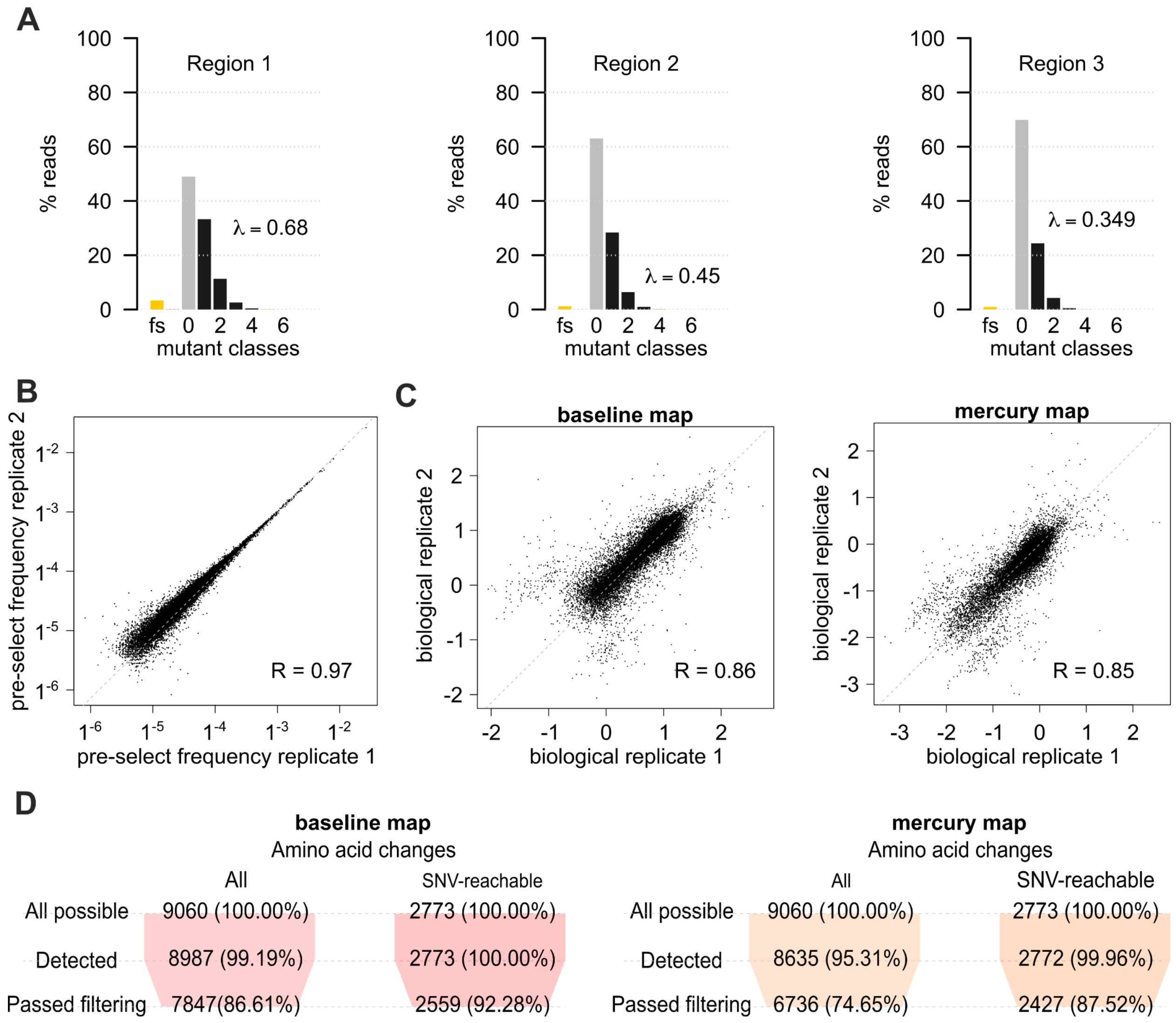


**Figure S2.** Characterization of the CPOX variant libraries used in generating the baseline and mercury maps. (A) Distribution of the number of missense variants in clones from the CPOX mutagenized libraries, and the fraction of clones carrying small indels resulting in frameshifts ("fs"). The average number of amino acid changes per clone (λ) is also estimated (see Methods). Correlation of variant frequencies between pre-selection libraries between (B) two separate transformed ('pre-select') pools and (C) between two post-selection biological replicates under both baseline (left) and mercury (right) environments. (D) Percentage of variants detected and passing quality control are shown for each CPOX map. For each map, the number of synonymous, nonsense, and missense substitutions (up to 19 possible) are shown across all residue positions, both before (left) and after (right) restricting to substitutions that are possible given a single nucleotide change. The three rows correspond to: 1) theoretically possible amino acid substitutions; 2) substitutions detected in the pre-selection condition; and 3) substitutions above a threshold pre-selection frequency.

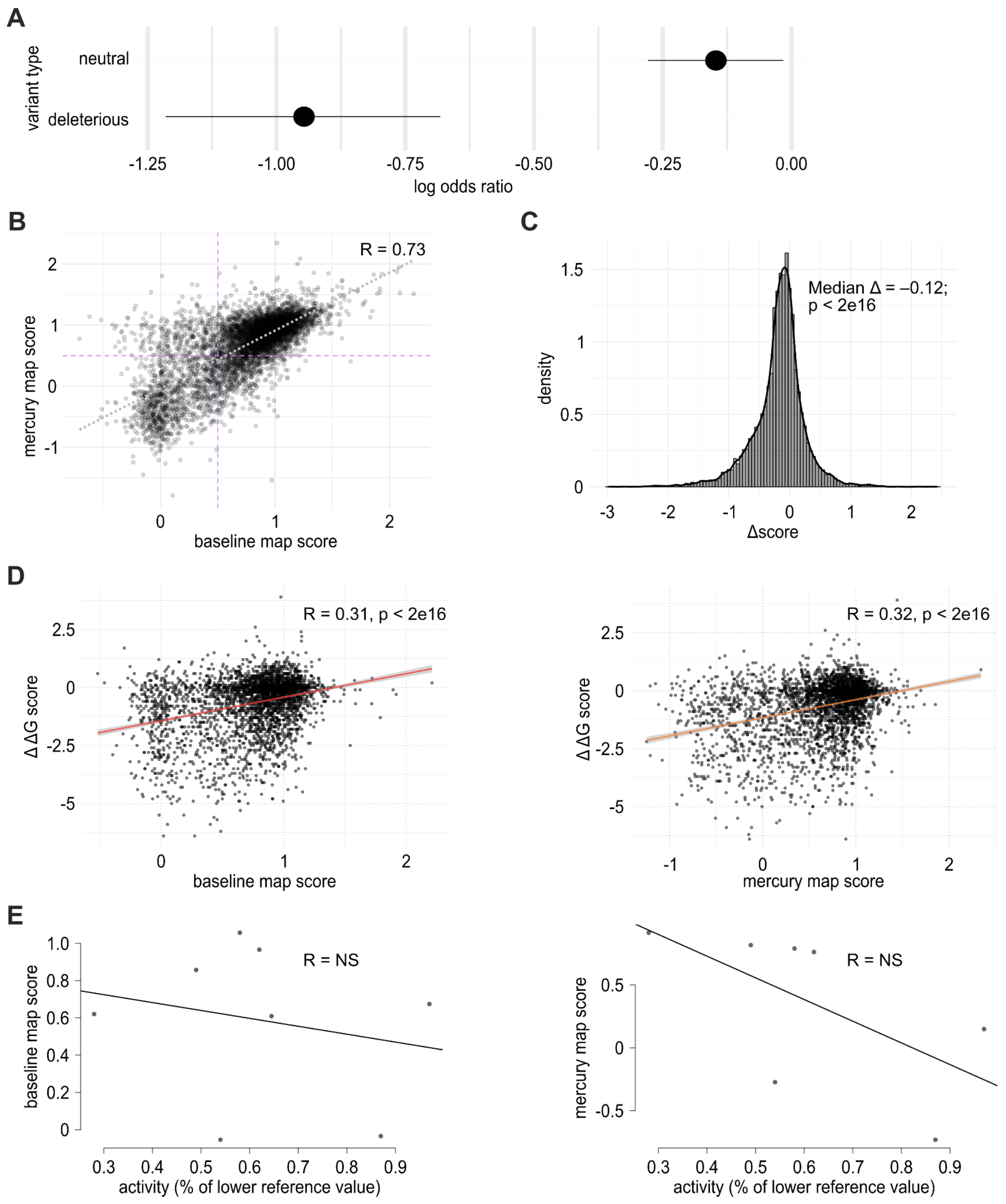


**Figure S3.** Quality control analysis of baseline and mercury map scores. (A) Log-odds ratios evaluating the depletion of missense variants with neutral or damaging baseline map scores in both UK Biobank and gnomAD population sequencing databases. Range lines represent 95% confidence intervals. (B) Correlation between baseline and mercury map scores for missense variants (Pearson's r = 0.73, $p <$ 2e-16). (C) Distribution of delta scores (mercury functional score − baseline functional score; Δmedian = 0.16, $p$ = 1e-3, Wilcoxon test; see Methods). (D) Correlation of positional median scores from each map with predicted folding free energy changes (Pearson's r = 0.31 and 0.32 for baseline and mercury maps, respectively; $p$ < 2e-16). (E) Correlation of positional median scores from each map with relative CPOX enzyme activity (variant activity divided by wild-type activity). NS, not significant

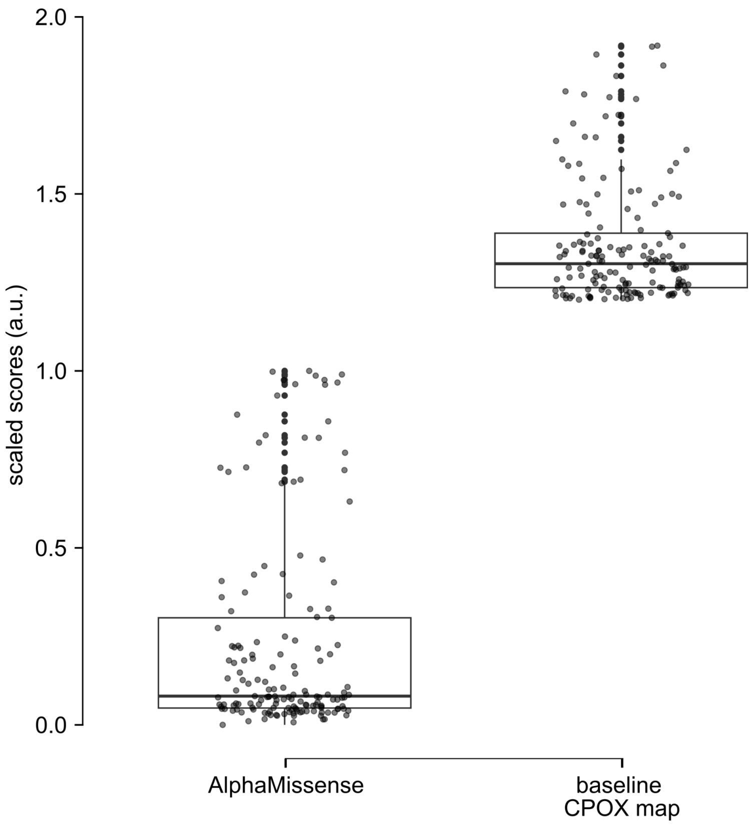


**Figure S4**. Comparison of ‘hyper-complementing’ scores from our baseline map and the computational predictor AlphaMissense (after transformation to a common scale). In this normalization, 0 represents null-like variants, scores near while 1 represents neutral variants, and scores significantly greater than 1 represent ‘hyper-complementing’ behavior which our phylogenetic analysis indicates (for this gene and assay) deleteriousness in humans. Boxes indicate interquartile range, with bold horizontal lines indicating medians. Whiskers indicate maxima and minima inside the 1.5x interquartile range.

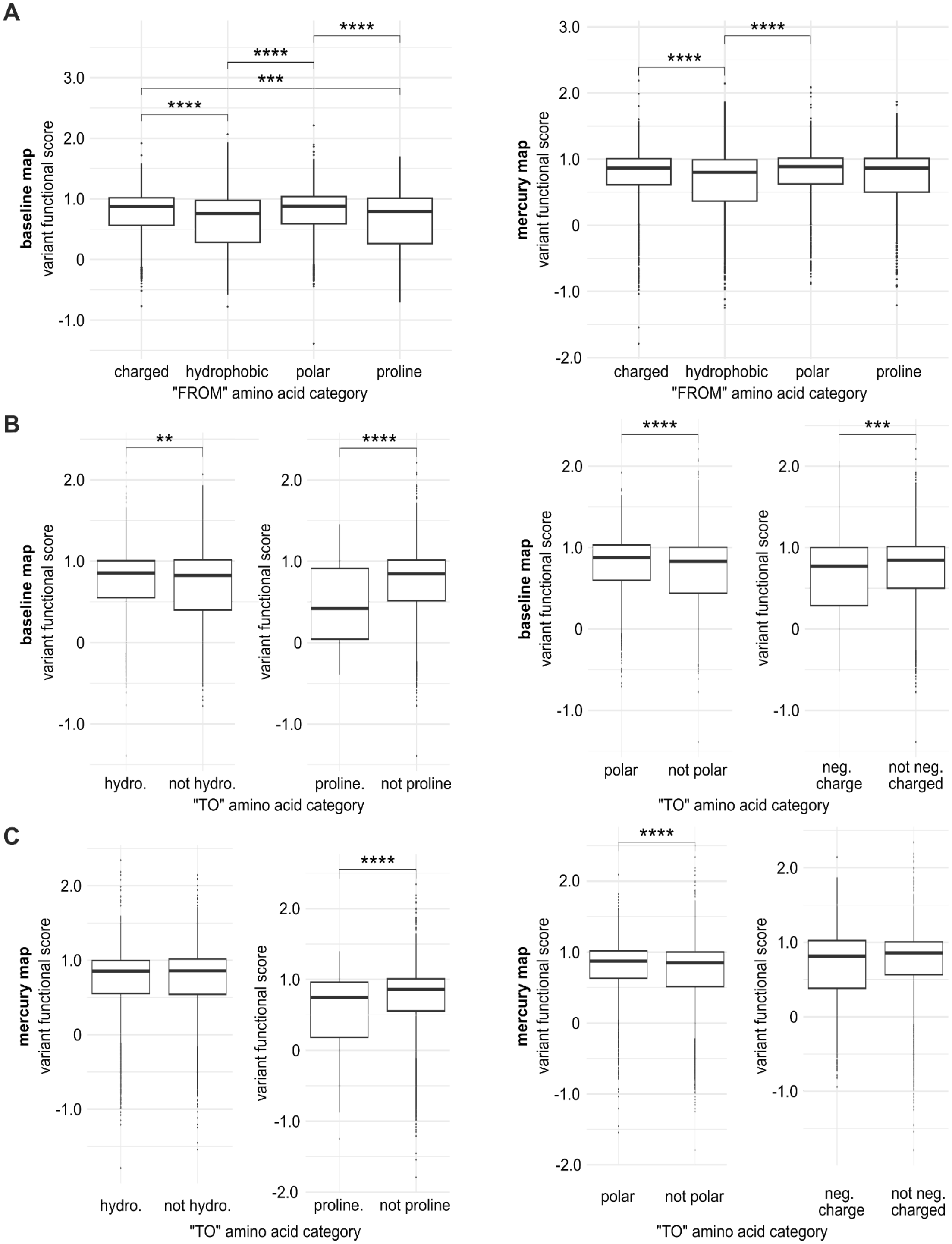


**Figure S5.** Distributions of variant functional scores by residue biochemical properties. (A) Distributions of variant functional scores for different initial (‘from’) amino acid categories for the (left) baseline map and (right) the mercury map. Scores for conservative substitutions compared to all other variants are shown for both the (B) baseline map and (C) mercury map in four subplots: hydrophobic vs. non-hydrophobic (left), proline vs. non-proline (mid-left), polar vs. non-polar (mid-right), and negative vs. non-negative (right). Bonferroni-adjusted Wilcoxon *p* values are indicated as follows: ****, $p \leq 0.0001$; ***, $p \leq 0.001$; **, $p \leq 0.01$; *, $p \leq 0.05$.

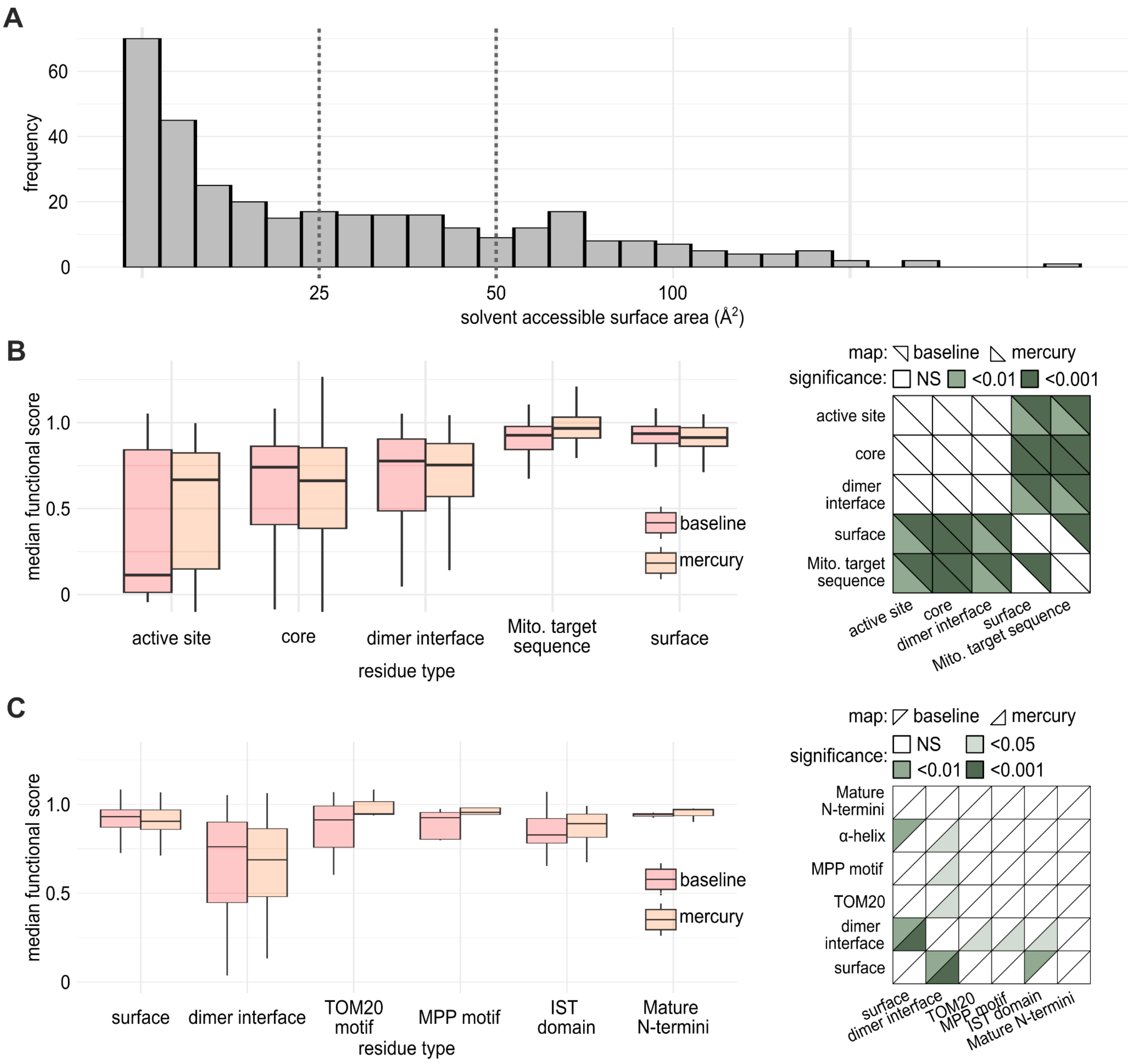


**Figure S6.** Modeling the effects of CPOX missense variants by residue and motif type. (A) Distribution of solvent accessible surface area (SASA) values for CPOX residues. Residues with SASA >50% were classified as exposed, whereas residues with SASA <25% were classified as buried (see Methods). (B) Left: For baseline and mercury maps, median functional scores of variants at positions corresponding to (1) active site residues required for oxidative decarboxylation of COPRO, (2) buried residues (<30% SASA), (3) residues at the dimerization interface, (4) residues in the mitochondrial targeting sequence (MTS), and (5) exposed residues (>50% SASA). Right: Triangular heatmap showing corresponding p-values (Wilcoxon tests) for residue categories in each map. (C) Left: For baseline and mercury maps, median functional scores of variants at positions corresponding to (1) exposed residues (>50% SASA), (2) residues at the dimerization interface, (3) TOM20 recognition motif, (4) mitochondrial processing peptidase (MPP) motif, (5) mitochondrial intermembrane space targeting (IST) domain, and (6) mature N-terminal residues. Right: Triangular heatmap showing corresponding *p*-values (Wilcoxon tests) for residue categories in each map.

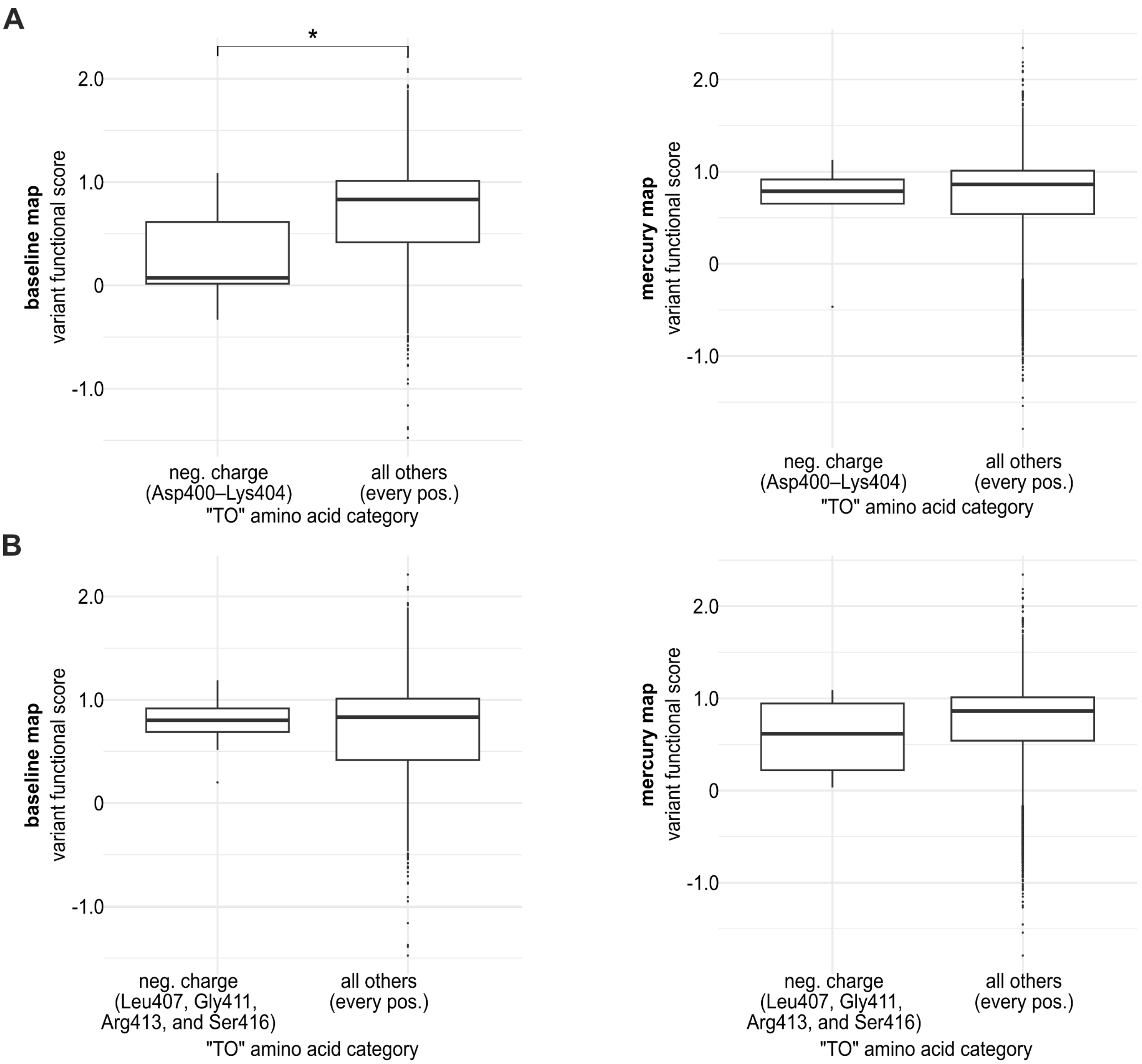


**Figure S7.** Distributions of functional scores for substitutions at the active-site regions of CPOX involved in substrate passage and the retention of harderoporphyrinogen. (A) Scores of substitutions to negatively-charged residues at Asp400–Lys404 (associated with harderoporphyrinogen retention) and (B) at Leu407, Gly411, Arg413, and Ser416 positions (lining the decarboxylation corridor), compared with all other variants across the baseline (left) and mercury (right) maps.

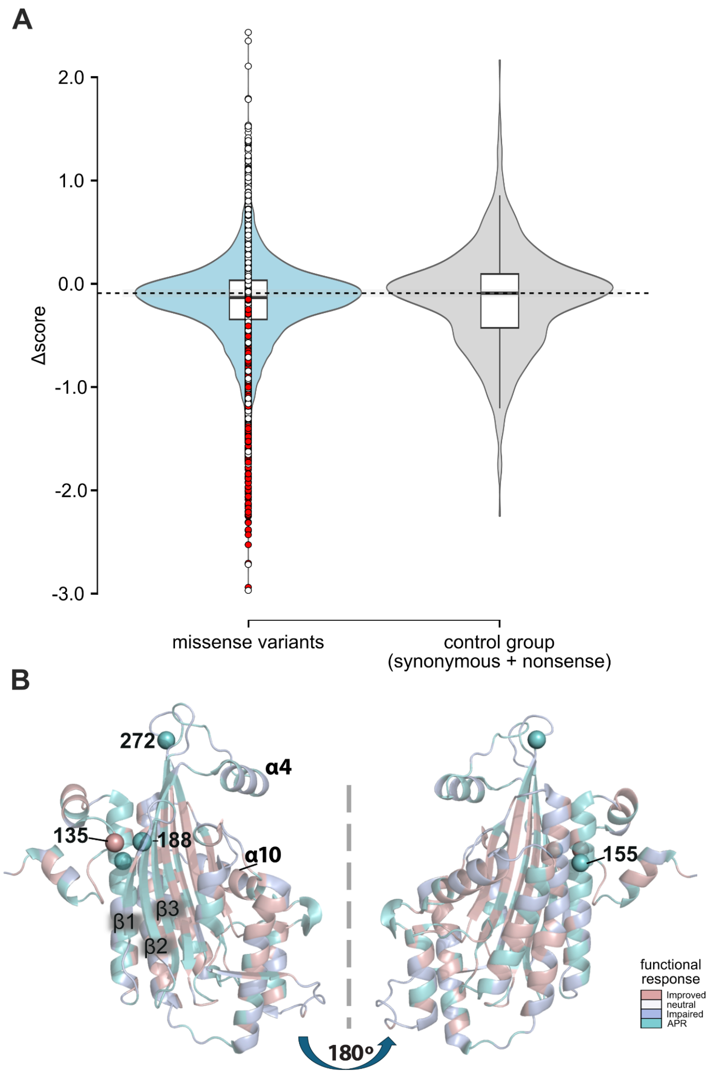


**Figure S8.** Identification and structural characterization of CPOX residues associated with an atypical porphyrinogenic response. (A) Distribution of delta scores ('Δscore'; mercury functional score − baseline functional score) for missense variants compared with synonymous and stop codon variants used as a null distribution. Missense variants whose 95% confidence intervals excluded the median Δscore of the control group (synonymous and stop variants) were classified as likely-having an atypical porphyrinogenic response (shown in red). Boxes indicate the interquartile range and the dotted line indicates the control group median. Whiskers indicate the minimum and maximum values. (B) Structure of CPOX colored by the median Δscore at each residue position, indicating functional response: improved (red; Δscore > 0), neutral (white; Δscore ≈ 0), atypical porphyrinogenic response (teal; Δscore < 0), and constitutively deleterious (blue; Δscore < 0 with a damaging baseline score).

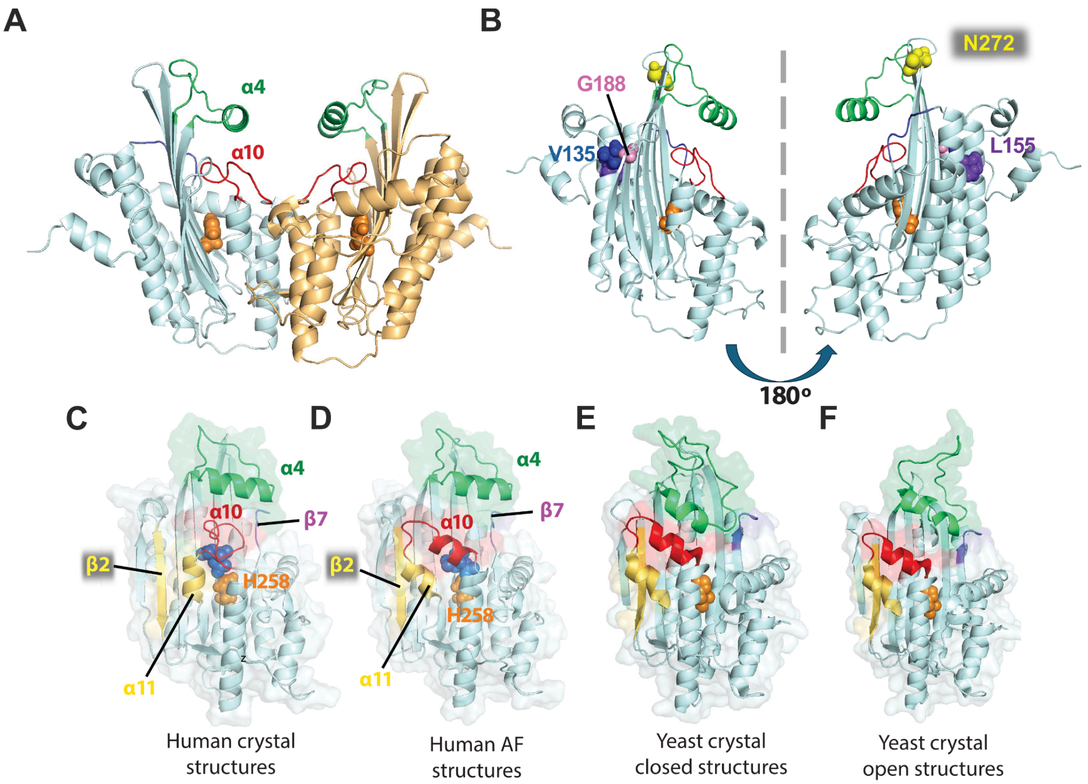


**Figure S9.** Comparison of CPOX crystal and AF model structures. **(A)** Homo-dimeric crystal structure of human CPOX (PDB ID:2AEX) showing the unstructured active site loop (red) and the lid (α4, green). The catalytic residue H258 is shown as orange spheres. **(B)** Residues (N272, L155, and G188) reflecting the missense variants used in our simulations are shown from two different views on a single monomer of the crystal structure. Single chains are shown with the interface at the front, comparing **(C)** human crystal structure, **(D)** human AF model and yeast Hem13 crystal structures in **(E)** closed (PDB id:1TLB) and **(F)** open states (PDB id: 1TKL). Secondary structural elements that regulate active site entrance are colored on the panels to display gate 1 (between α4 and active site loop), gate 2 (active site loop and β7) and gate 3 (β2 and α11). The overlayed surface and cartoon representation displays significant differences in gate 1. Open gate 1 exists in the human CPOX AF model and the open state of the yeast Hem13 structure. Note that the active site loop (red) indicated by α10 consists of only a single turn in the human crystal structure.

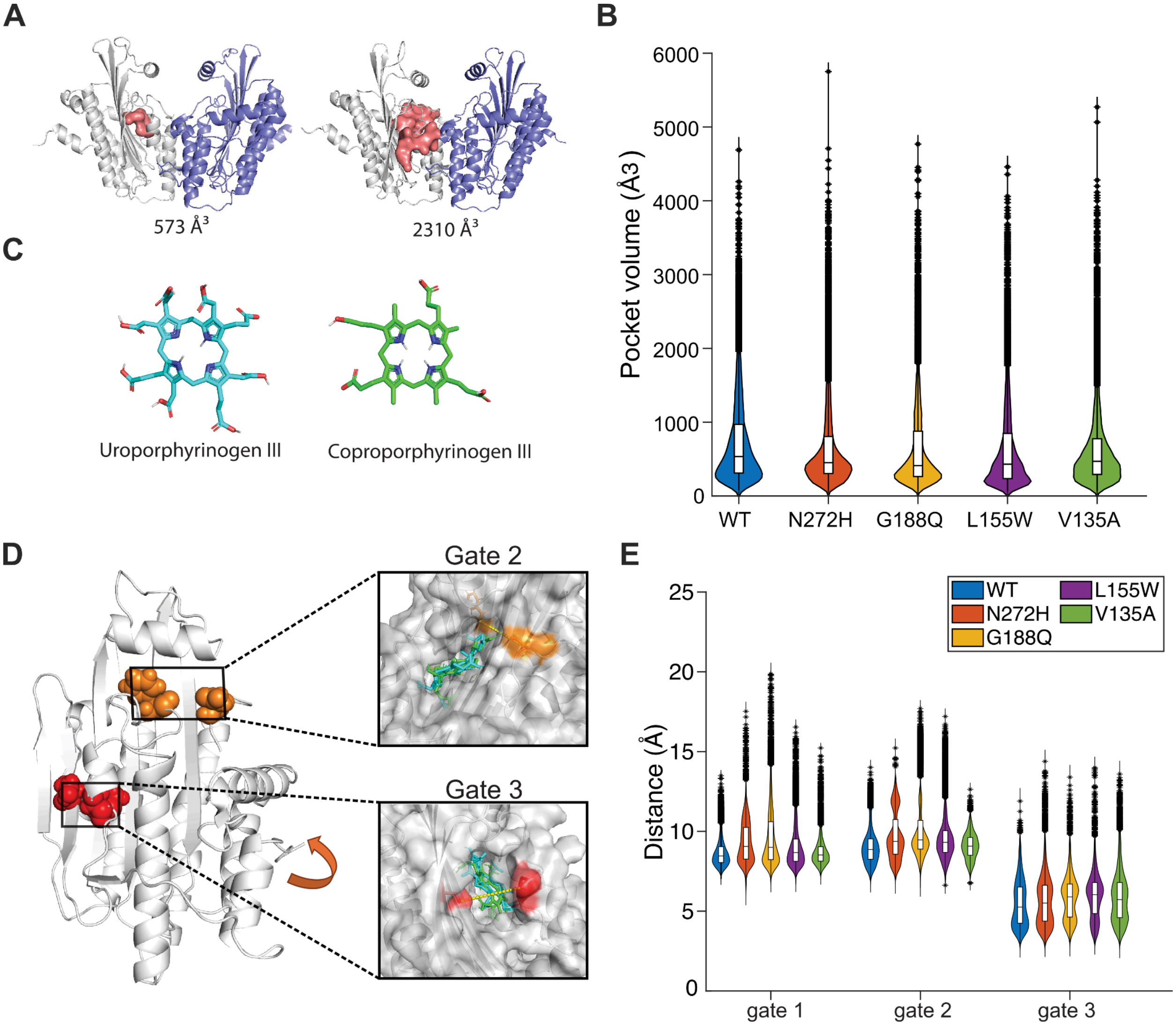


**Figure S10.** Features of active site entrances/gates and pocket volumes in the presence of the unstructured loop. (**A**) Dynamics of the active site pocket, illustrating the volumes of the initial (left, 573 Å³) and open (right, 2310 Å³) pocket conformations. (**B**) Active site pocket volume distributions for WT and variants based on simulations based on the human CPOX crystal structure. **(C)** Chemical structures of COPRO and URO used for molecular docking. COPRO (CID 321) and URO (HMDB0001086) were retrieved from PubChem CID 321[13], and the Human Metabolome Database [14], respectively. (**D**) Gates 2 and 3 are used by the substrates as entry points to the active site pocket when the loop is unstructured. (**E**) Distance distributions for each gate across simulations for WT and variants. Gate 1 remains closed due to the unstructured loop.

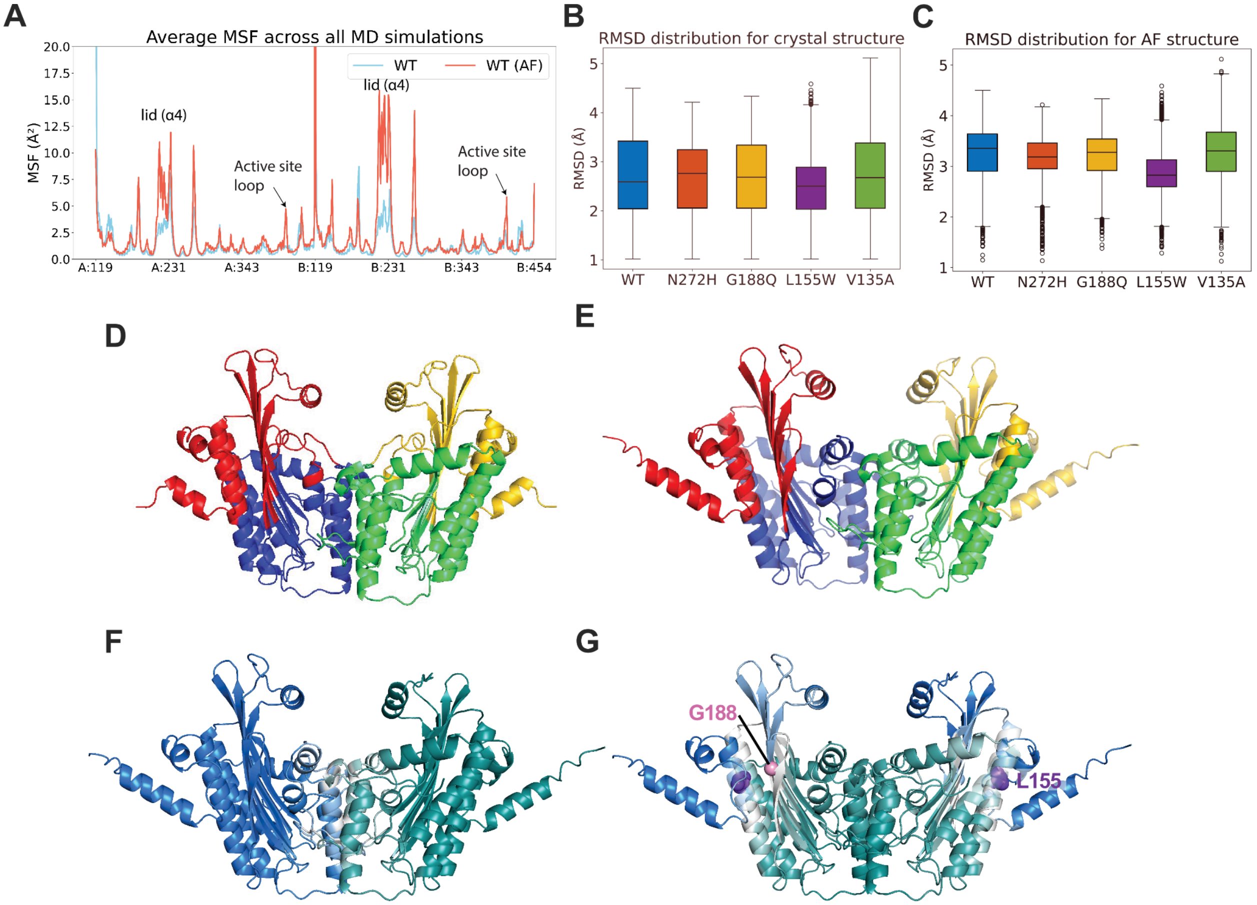


**Figure S11.** Comparison of CPOX dynamics between structures where the active site loop is unstructured (crystal) vs. helical (AF). (**A**) Mean-square fluctuations from WT simulations indicate higher mobility of lid and active site loop for helical loop (red) in comparison to the unstructured loop (blue). Distribution of root-mean-square deviation (RMSD) for (**B**) helical loop and (**C**) unstructured simulations. Dynamic domains (identified by the SPECTRUS server) reveal distinct coupling of the active site loop in (**D**) crystal structure and (**E**) AF model. (**F**) GNM global mode 1 for AF model indicates anticorrelated motion of the two chains (blue and green) with the hinge residues located at the dimer interface (white). (**G**) GNM mode 2 further divides each chain into two anti-correlated domains, i.e. the lid and the base being separated by flexible/hinge residues (white). Notably, L155 (purple) and G188 (pink) act as hinge points in this mode.

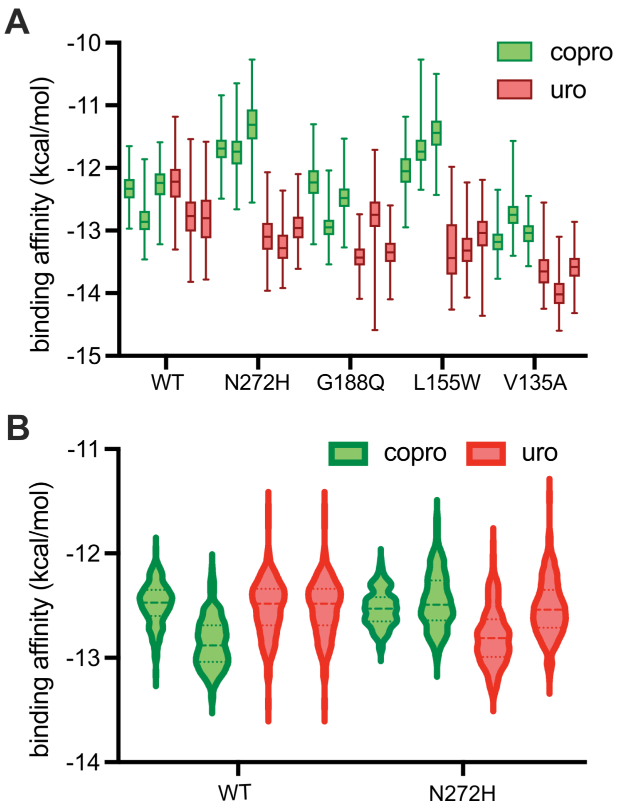


**Figure S12.** Distribution of binding affinities (ΔG, PRODIGY-LIG)[10] across the MD frames for the specific chain (A or B) with the best docking score. (A) AF simulations for WT and variant models (N272H, G188Q, L155W) are shown using Box plots based on three independent runs for each ligand. (B) Simulations for WT and N272H complexes based on the CPOX crystal structure are shown via violin plots based on two independent runs (median dashed, quartiles dotted) are shown for COPRO and URO.

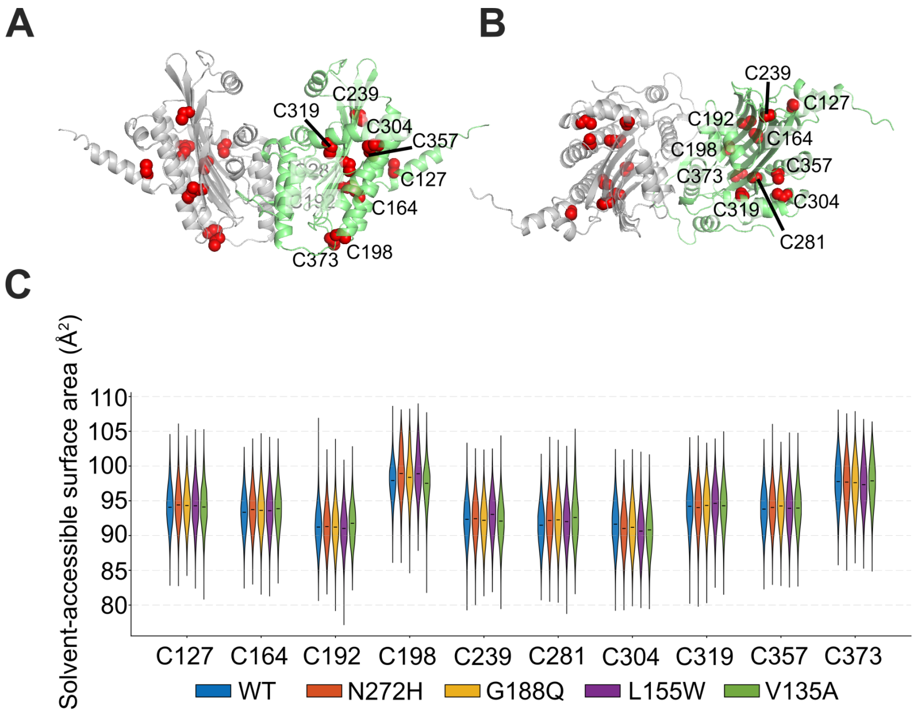


**Figure S13.** Solvent accessibility of cysteines as mercury binding sites in MD simulations. Cysteines (*red spheres*) are shown on the (**A**) top and (**B**) side views of the AF2 dimer structure. (**C**) SASA distributions (violin plots) for these residues are plotted using snapshots from AF simulations. There are no significant differences in SASA distributions between WT and variants, which suggests comparable effects due to mercury-binding[15].

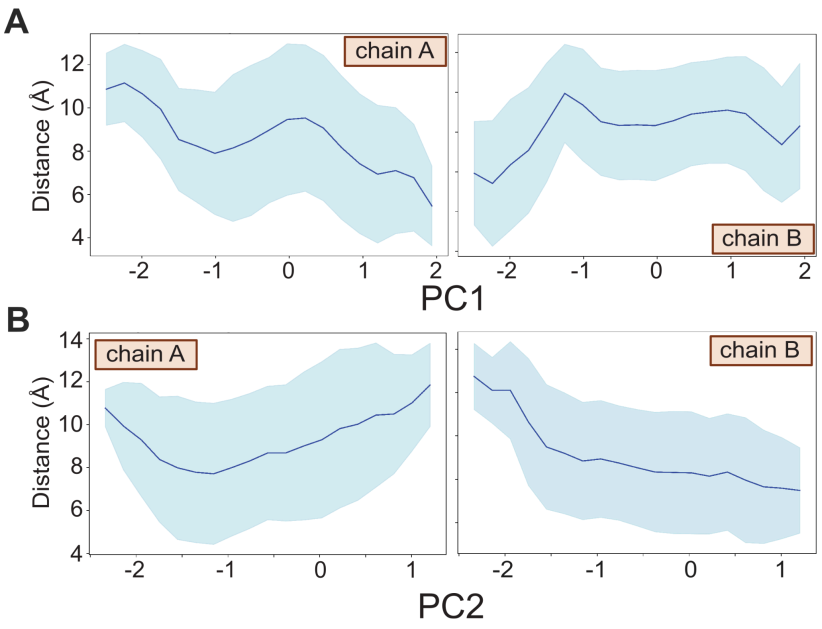


**Figure S14.** Relationship between Gate 2 opening size and the progression along **(A)** PC1 and **(B)** PC2 for each chain.

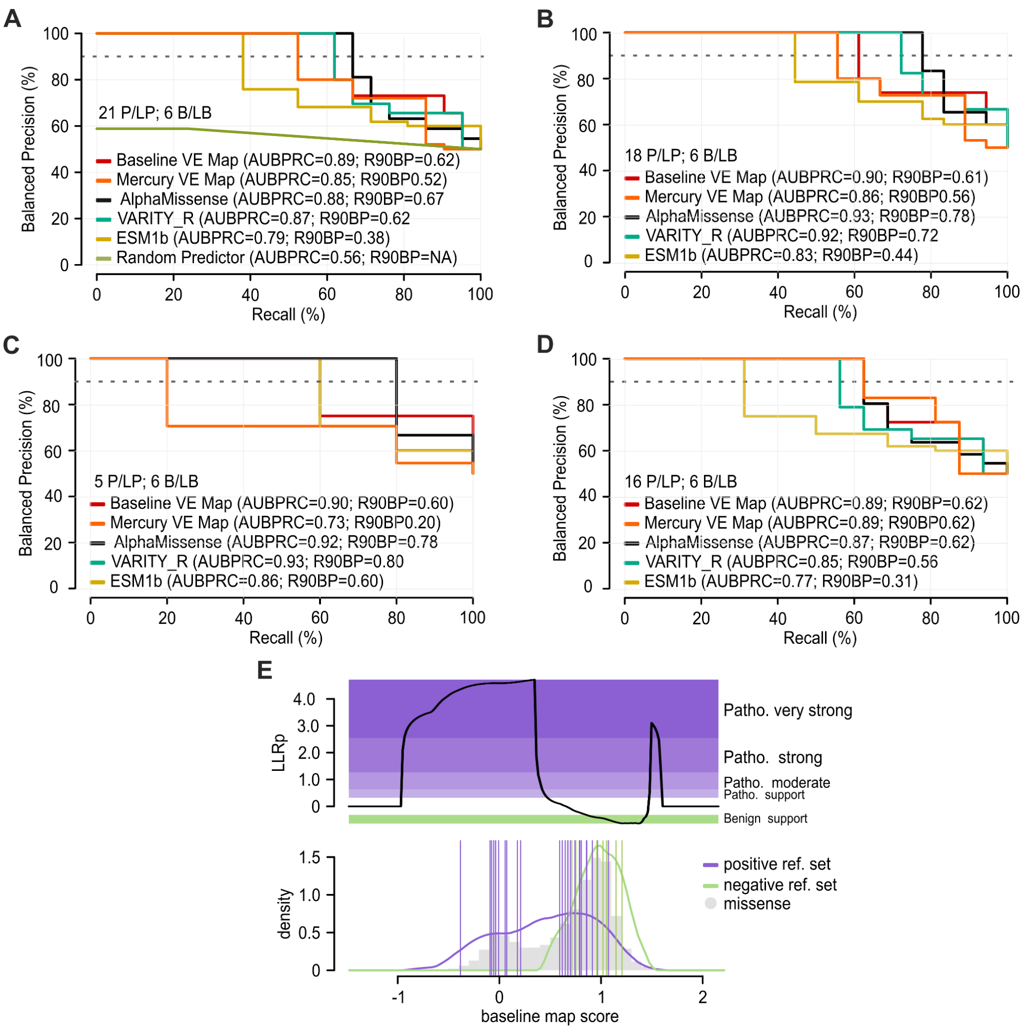


**Figure S15.** Performance of both CPOX variant maps and computational predictors in distinguishing positive and negative reference variants, and evidentiary value of baseline map scores for clinical variant interpretation. Here we evaluate precision (fraction of variants scoring below each threshold functional score that are in the positive reference set containing pathogenic variants) vs recall (fraction of positive reference variants with functional scores below threshold). Precision has been transformed to reflect performance in a balanced test setting where positive and negative sets contain the same number of variants. Balanced precision–recall curves are shown for the baseline map (red), mercury map (orange), a random predictor (lime green), and computational predictors: AlphaMissense (black), VARITY_R (turquoise), and ESM1b (gold). Performance was evaluated using (A) all reference variants, (B) variants excluding positions 1–110 corresponding to the mitochondrial targeting sequence, and reference sets defined only with (C) or without (D) harderoporphyria (HP)-associated pathogenic variants, and summarized by the area under the balanced precision–recall curve (AUBPRC) and recall at 90% balanced precision (R90BP). Positive and negative reference set sizes (P/LP and B/PB, respectively; see Methods) are indicated. (E) Transformation functions represent variant effects in terms of the strength of evidence for and against pathogenicity, that is, a log-likelihood ratio (LLR) of pathogenicity. The functions (top) express the log ratio between the likelihood of observing a given score in the score distribution of the positive reference (purple; total set) set as opposed to that of the negative reference set (green). Gray histogram bars show the distribution of missense variants for comparison.

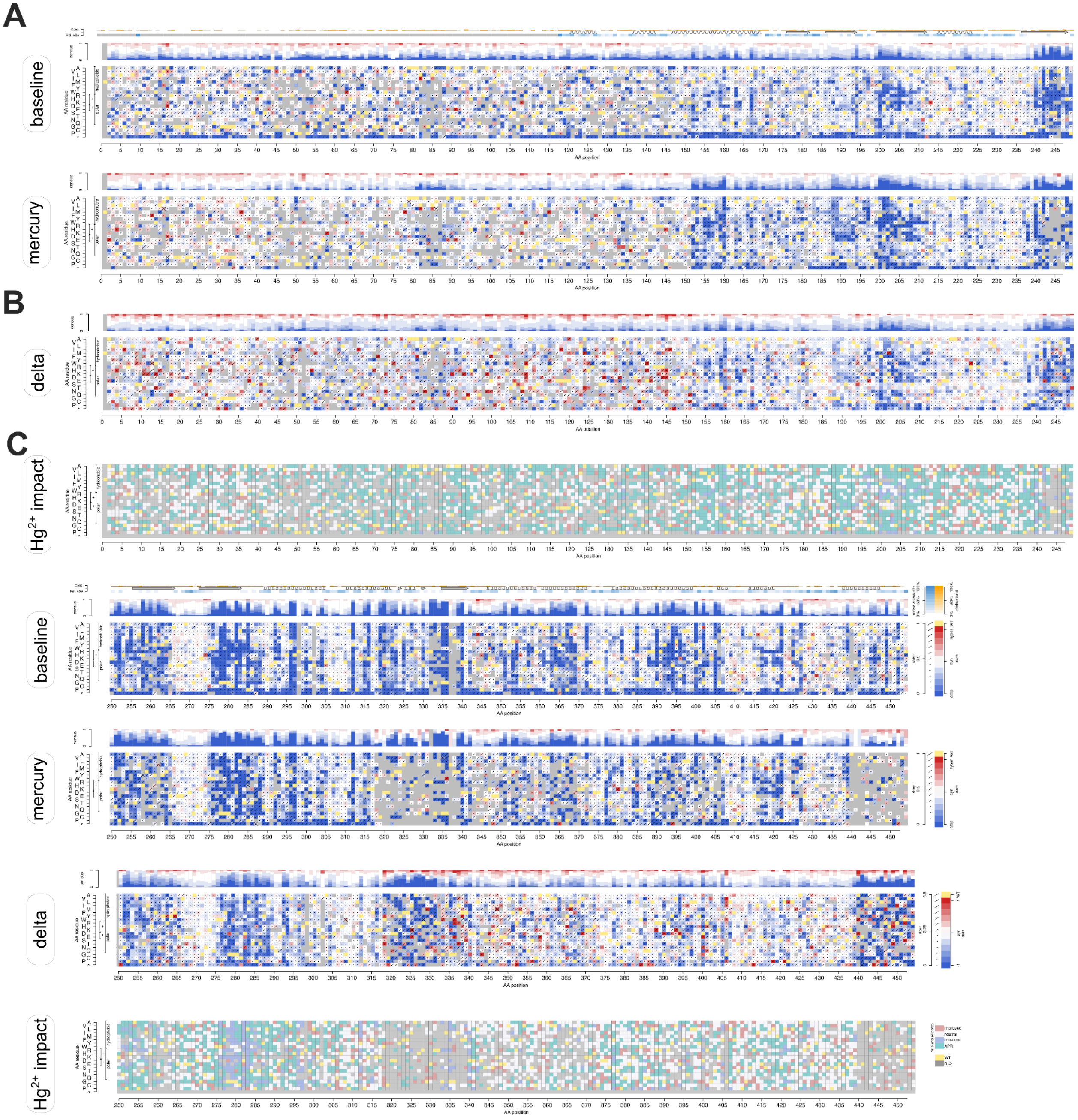


**Figure S16.** Full-sized CPOX variant effect maps. (A) Complete functional maps for baseline and mercury. Colors and labels are as in Figure 2. (B) Delta landscape (unscaled mercury functional score − baseline functional score) for missense variants. (C) Map of mercury ($Hg^{2+}$) impact. Substitutions were coloured by functional response: improved (red; Δscore > 0), neutral (white; Δscore ≈ 0), atypical porphyrinogenic response (teal; Δscore < 0), and constitutively deleterious (blue; Δscore < 0 with a damaging baseline score).

**Supplementary Tables**

| **Model** | **ΔAIC relative to best model** |
|---|---|
| Penalize enhancing mutations | 0 |
| Cap score of enhancing mutations at WT level | 197.8 |
| Enhancing mutations are beneficial | 199.4 |

**Table S1.** Comparing different models for effects of activity enhancing mutations.

| **Initial structure** | **State** | **MD details** | **WT** | **N272H** | **G188Q** | **L155W** | **V135A** |
|---|---|---|---|---|---|---|---|
| Crystal structure (2aex) | apo | # of runs | 5 | 5 | 5 | 5 | 5 |
| | | Run duration (ns) | 100 | 200 | 100 | 100 | 100 |
| | | Total (ns) | 500 | 1000 | 500 | 500 | 500 |
| AF model | apo | # of runs | 2 | 2 | 2 | 2 | 2 |
| | | Run duration (ns) | 200 | 200 | 200 | 200 | 200 |
| | | Total (ns) | 400 | 400 | 400 | 400 | 400 |
| AF model | docked complex | # of runs | 3 | 3 | 3 | 3 | 3 |
| | | Run duration (ns) | 200 | 200 | 200 | 200 | 200 |
| | | Total (ns) | 600 | 600 | 600 | 600 | 600 |
| Crystal structure (2aex) | docked complex | # of runs | 2 | 2 | - | - | - |
| | | Run duration (ns) | 200 | 200 | - | - | - |
| | | Total (ns) | 400 | 400 | - | - | - |

**Table S2.** Details of MD simulations for WT and variants

| Crystal Mutation | Binding affinity for best pose (kcal/mol) | | | Active site occupancy (%) | | Total no of docking poses | Binding affinity from PRODIGY-LIG (kcal/mol) | |
|---|---|---|---|---|---|---|---|---|
| | COPRO | URO | Δ(COPRO - URO) | COPRO | URO | | COPRO | URO |
| WT | **-9.6** | -7.9 | -1.7 | **10** | 5 | 200 | -12.7 ± 0.3 | -12.5 ± 0.3 |
| N272H | -10.0 | **-10.2** | 0.2 | 24 | **69** | 300 | -12.5 ± 0.2 | -12.7 ± 0.3 |
| L155W | -8.4 | **-10.3** | 1.9 | 3 | **20** | 200 | - | - |
| G188Q | -8.8 | **-9.1** | 0.3 | 7 | **22** | 100 | - | - |

**Table S3.** Substrate docking using snapshots with unstructured active site loop (crystal simulations)

| **Model** | **Gate 1** | | TH (Å) | **Gate 2** | | TH (Å) | **Gate 3** | | TH (Å) |
|---|---|---|---|---|---|---|---|---|---|
| | Atom1 | Atom 2 | | Atom 1 | Atom 2 | | Atom 1 | Atom 2 | |
| Crystal | Q221 (CA) | F405 (CA) | 12 | D341 (CA) | F408 (CA) | 10 | S191 (OG) | M419 (CE) | 10 |
| AF [c] | Q221 (CA) | R401 (CA) | 15 | D340 (OD2) | R401 (NH2) | 10 | S191 (OG) | M419 (CE) | 10 |

[a] For each gate, the residue pairs that are used for distance calculations are provided together with the corresponding atoms (in PDB format).
[b] The threshold (TH) value is used to assess putative gate opening. Note that a higher threshold for gate 1 was chosen for AF runs as this gate stays mostly open. In contrast the active site pockets and gates were comparatively smaller due to unstructured loop.
[c] As explained in text, gate opening for the AF snapshots is monitored by (α10 - α4) distance for gate 1, (α10 - β7) distance for gate 2 and (β2 - α11) distance for gate 3 . In the AF model, the active site loop contains α10, whereas in the crystal structure this helix has a single turn that completely unfolds during simulations. Different atom pairs are used for AF and crystal simulations for gates 1 and 2.

**Table S4.** Active site gates[a] and corresponding minimum distances[b] for gate opening

### REFERENCES


1. van Loggerenberg, W. *et al.* Systematically testing human HMBS missense variants to reveal mechanism and pathogenic variation. *Am. J. Hum. Genet.* **110**, 1769–1786 (2023).
2. Weile, J. *et al.* A framework for exhaustively mapping functional missense variants. *Mol. Syst. Biol.* **13**, 957 (2017).
3. Bloom, J. D. An experimentally determined evolutionary model dramatically improves phylogenetic fit. *Mol. Biol. Evol.* **31**, 1956–1978 (2014).
4. Bloom, J. D. Identification of positive selection in genes is greatly improved by using experimentally informed site-specific models. *Biol. Direct* **12**, 1 (2017).
5. Cheng, J. *et al.* Accurate proteome-wide missense variant effect prediction with AlphaMissense. *Science* **381**, eadg7492 (2023).
6. Lee, D.-S. *et al.* Structural basis of hereditary coproporphyria. *Proc. Natl. Acad. Sci. U. S. A.* **102**, 14232–14237 (2005).
7. Phillips, J. D. *et al.* Crystal structure of the oxygen-dependant coproporphyrinogen oxidase (Hem13p) of Saccharomyces cerevisiae. *J. Biol. Chem.* **279**, 38960–38968 (2004).
8. Silva, P. J. & Ramos, M. J. Computational characterization of the substrate-binding mode in coproporphyrinogen III oxidase. *J. Phys. Chem. B* **115**, 1903–1910 (2011).
9. Ponzoni, L., Polles, G., Carnevale, V. & Micheletti, C. SPECTRUS: A dimensionality reduction approach for identifying dynamical domains in protein complexes from limited structural datasets. *Structure* **23**, 1516–1525 (2015).
10. Vangone, A. *et al.* Large-scale prediction of binding affinity in protein-small ligand complexes: the PRODIGY-LIG web server. *Bioinformatics* **35**, 1585–1587 (2019).
11. Li, T. & Woods, J. S. Cloning, expression, and biochemical properties of CPOX4, a genetic variant of coproporphyrinogen oxidase that affects susceptibility to mercury toxicity in humans. *Toxicol. Sci.* **109**, 228–236 (2009).
12. Li, T. & Woods, J. S. Cloning, expression, and biochemical properties of CPOX4, a genetic variant of coproporphyrinogen oxidase that affects susceptibility to mercury toxicity in humans. *Toxicol. Sci.* **109**, 228–236 (2009).
13. Kim, S. *et al.* PubChem 2025 update. *Nucleic Acids Res.* **53**, D1516–D1525 (2025).
14. Wishart, D. S. *et al.* HMDB 5.0: The Human Metabolome Database for 2022. *Nucleic Acids Res.* **50**, D622–D631 (2022).
15. Lu, C.-H. *et al.* MIB2: metal ion-binding site prediction and modeling server. *Bioinformatics* **38**, 4428–4429 (2022).
